# Competition for shared resources increases dependence on initial population size during coalescence of gut microbial communities

**DOI:** 10.1101/2023.11.29.569120

**Authors:** Doran A. Goldman, Katherine S. Xue, Autumn B. Parrott, Rashi R. Jeeda, Lauryn R. Franzese, Jaime G. Lopez, Jean C. C. Vila, Dmitri A. Petrov, Benjamin H. Good, David A. Relman, Kerwyn Casey Huang

## Abstract

The long-term success of introduced populations depends on their initial size and ability to compete against existing residents, but it remains unclear how these factors collectively shape colonization. Here, we investigate how initial population (propagule) size and resource competition interact during community coalescence by systematically mixing eight pairs of *in vitro* microbial communities at ratios that vary over six orders of magnitude, and we compare our results to a neutral ecological model. Although the composition of the resulting co-cultures deviated substantially from neutral expectations, each co-culture contained species whose relative abundance depended on propagule size even after ∼40 generations of growth. Using a consumer-resource model, we show that this dose-dependent colonization can arise when resident and introduced species have high niche overlap and consume shared resources at similar rates. This model predicts that propagule size will have larger, longer-lasting effects in diverse communities in which niche overlap is higher, and we experimentally confirm that strain isolates show stronger dose dependence when introduced into diverse communities than in pairwise co-culture. This work shows how neutral-like colonization dynamics can emerge from non-neutral resource competition and have lasting effects on the outcomes of community coalescence.

## Introduction

Colonization by new species plays a major role in shaping community composition and function, but the outcomes of species introductions are notoriously difficult to predict^1–7^. To colonize, a species must disperse from its original population^8–10^, overcome the stochastic effects of ecological drift^11,12^, and compete successfully against resident species for a niche in the community^13–17^. Models of community assembly place different emphasis on the role of neutral forces—namely, dispersal and ecological drift—compared to competition^15,18,19^, and it remains unclear how these neutral and non-neutral forces interact to shape colonization outcomes.

Ecological models differ especially in their predictions about how colonization outcomes depend on the size and frequency of introduced populations, together known as propagule pressure^5,20^. The neutral theory of biodiversity, which assumes that species are ecologically equivalent, predicts that the abundance of an introduced species will remain proportional to its initial propagule size^15,18,21^. In contrast, in classical consumer-resource models, each introduced species reaches an equilibrium abundance determined by its competitive ability, regardless of its initial abundance^19,22^. Investigating the effect of propagule size therefore provides an opportunity to quantify the impact of neutral and non-neutral forces on colonization.

So far, empirical studies have come to contradictory conclusions about the effect of propagule size on colonization. Large propagule size is often associated with successful colonization^23–27^, consistent with neutral expectations. However, there are many cases in which large propagules fail to colonize or small propagules successfully establish^5,20,21,28–34^, showing that propagule size is not always the main determinant of colonization success. Many of these studies focus on the introduction of a small number of species^24–26,28,31,32,34^, making it difficult to generalize their conclusions across species and community contexts. Resolving these discrepancies requires systematic, large-scale quantification of the effect of propagule size.

To address this gap, we investigated the impact of propagule size on colonization during coalescence of *in vitro* gut microbial communities. Understanding the effect of propagule size in the human gut is important for designing microbiome therapeutics^35,36^, but prior studies have come to inconsistent conclusions about how inoculation dose affects the outcome of probiotic introductions and fecal microbiota transplants^37–42^. To investigate propagule size in a more tractable, laboratory setting, we used stool-derived *in vitro* communities, which are stable, diverse, reproducible models of the gut microbiome that can recapitulate *in vivo* responses to perturbation^22,43–47^. We mixed pairs of *in vitro* communities at ratios that varied over six orders of magnitude, allowing us to observe the outcomes of hundreds of species introductions at once. By combining these experiments with consumer-resource modeling, we show how resource competition can amplify the effects of initial population size in complex communities, with lasting effects on the outcomes of community coalescence^16,48^.

## Results

### Diversity and composition of co-culture communities vary across mixture ratios but deviate from neutral predictions

We performed coalescence experiments *in vitro* using a set of diverse, stable communities of gut microbes. Following a previously established protocol^46^, we derived these communities from stool samples of eight healthy human subjects and passaged them 15 times with a 1:200 dilution in fresh modified Brain Heart Infusion + mucin (mBHI+mucin) medium every 48 h (**Fig. 1a, Methods**). To assess community composition, we performed 16S rRNA gene amplicon sequencing on multiple passages from each community and tracked the relative abundances of amplicon sequence variants (ASVs, a rough taxonomic equivalent of species). Community diversity decreased in the first few passages after initial laboratory inoculation (**Fig. 1b, Extended Data Fig. 1a,b**), likely because some taxa in the inoculum were nonviable or were outcompeted during *in vitro* passaging. Nonetheless, diversity stabilized after ∼3 passages, and most communities thereafter contained ASVs that comprised >50% of the relative abundance of the stool inoculum (**Extended Data Fig. 1c-f**). Thus, these stool-derived *in vitro* communities are diverse and stable models of stool microbiotas that can be used to study ecological interactions among gut microbes.

**Figure 1:**
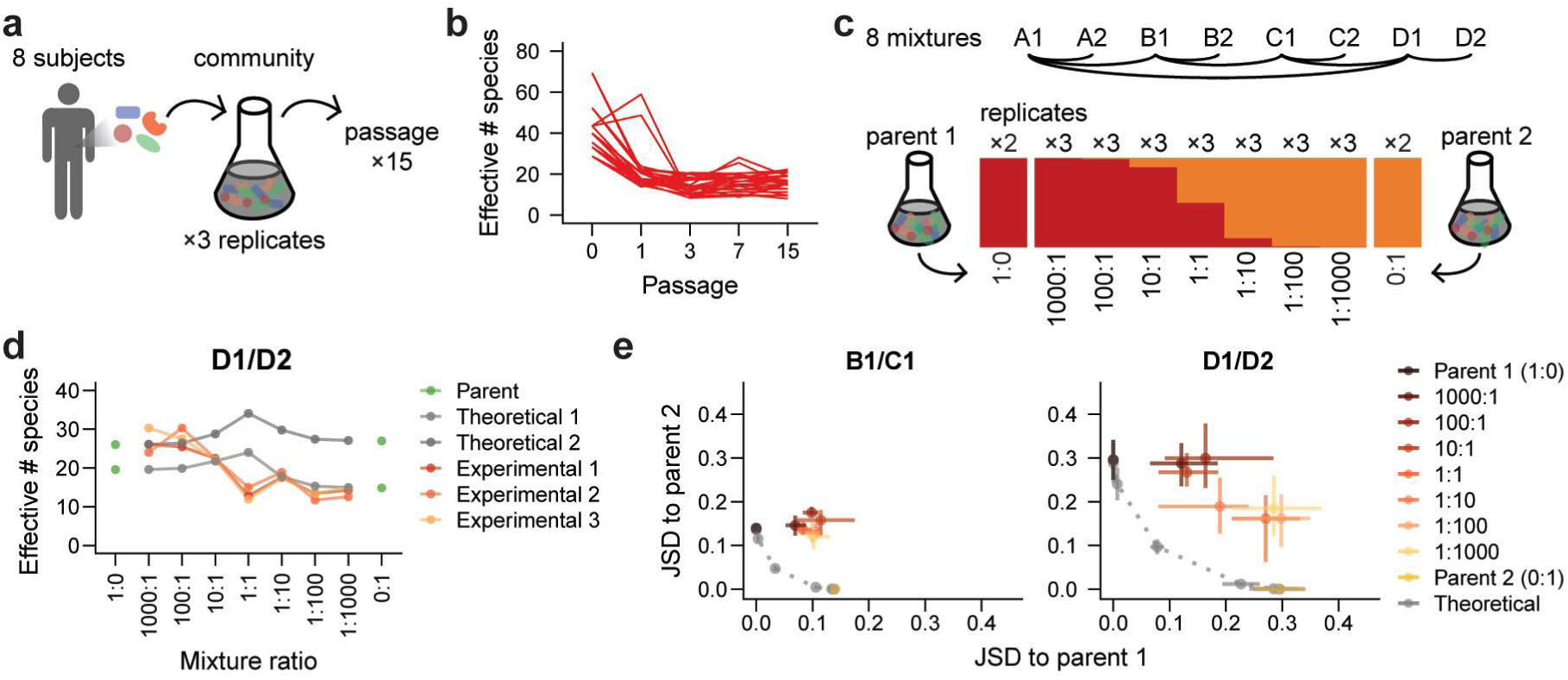
Diversity and composition of co-cultures vary by mixture ratio and deviate from neutral predictions. a) *In vitro* communities were inoculated in triplicate from stool samples collected from eight healthy human subjects and passaged 15 times to reach stability. b) The diversity of each community over time, quantified as the effective number of species (*e*^H′^) calculated from the Shannon diversity index (*H*′), initially decreased and then plateaued by passage 3. c) Experimental design: eight pairs of *in vitro* communities were mixed in triplicate at mixture ratios ranging from 1000:1 to 1:1000. The resulting co-cultures were passaged five times. d) The diversity of co-culture communities was similar to or lower than the diversity of parent communities across mixture ratios. Gray lines represent both replicates of predicted neutral mixtures for an example mixture based on the diversity of replicate parent communities (1:0 and 0:1, shown in green), and orange lines represent three replicates of experimental community mixtures. Diversity was quantified by the effective number of species (*e*^H′^) calculated from the Shannon diversity index (*H′*). e) Variation of community composition across example mixtures and mixture ratios. Each plot shows the Jensen-Shannon divergence (JSD) of all co-culture and theoretical communities relative to both parent communities (brown and gold) for a single mixture. Grey points along the dotted line represent theoretical mixtures, and colored points represent data from experimental co-culture communities. Points show averages and error bars show full range of values across inoculation replicates.

To systematically assess the effects of initial propagule size on colonization, we mixed eight pairs of *in vitro* “parent” communities at seven mixture ratios ranging from 1000:1 to 1:1000 (**Fig. 1c, Extended Data Fig. 2, Table S1**). We co-cultured mixtures in triplicate for five passages to allow composition to re-equilibrate, and we passaged two replicates of each parent community alongside to allow for comparison. To assess the impact of propagule size on community composition and the colonization of individual species, we compared our data to the predictions of ecological neutral theory^15,18,21^. We computationally generated a set of theoretical co-cultures whose composition was determined by the propagule size of each species in the initial mixture (**Extended Data Fig. 3a-c**), and we quantified the similarity of our data to these theoretical communities.

Under neutral expectations, co-culture diversity should be maximal at the 1:1 mixture ratio. However, we found that experimental co-cultures consistently maintained diversities similar to or lower than one of the parent communities (**Fig. 1d, Extended Data Fig. 4**). This finding suggests that non-neutral competition occurs during community coalescence, preventing the neutral coexistence of all species. We also compared each experimental co-culture to its corresponding theoretical co-culture using the Jensen-Shannon divergence (JSD, **Methods**). Co-cultures consistently differed from the corresponding neutral theoretical expectation at every mixture ratio (**Fig. 1e, Extended Data Fig. 5**), as we observed with community diversity. Despite these deviations from neutral predictions, the composition of many, though not all, co-cultures varied substantially across mixture ratios (**Fig. 1e, Extended Data Fig. 5**), highlighting the continued impact of propagule size after ∼40 generations of growth. These patterns of composition suggest that neither neutral theory nor consumer-resource models alone can fully predict the varied outcomes of community coalescence.

### Dose-dependent colonizers are present in every community mixture

To investigate how the behaviors of individual species give rise to non-neutral but propagule size-dependent patterns of community composition, we examined colonization outcomes at the ASV level. We compared the relative abundance of each ASV in experimental co-cultures after five passages to the neutral theoretical prediction determined solely by its propagule size in the initial mixture (**Extended Data Fig. 6**). In contrast to neutral predictions, many ASVs that started at higher abundance in one parent community maintained a consistently high or low relative abundance across mixture ratios in experimental co-cultures (*n*=32 (7.8%) of the 410 ASVs above the limit of detection, **Methods**; **Fig. 2a, Extended Data Fig. 6, 7a,b**). We classified these dose-independent ASVs as strong or weak colonizers depending on whether their experimental relative abundance was greater or less than the neutral prediction, respectively. For these strong and weak colonizers, deviations from neutral predictions likely reflect competitive advantages or disadvantages that ultimately outweighed the effects of propagule size.

**Figure 2:**
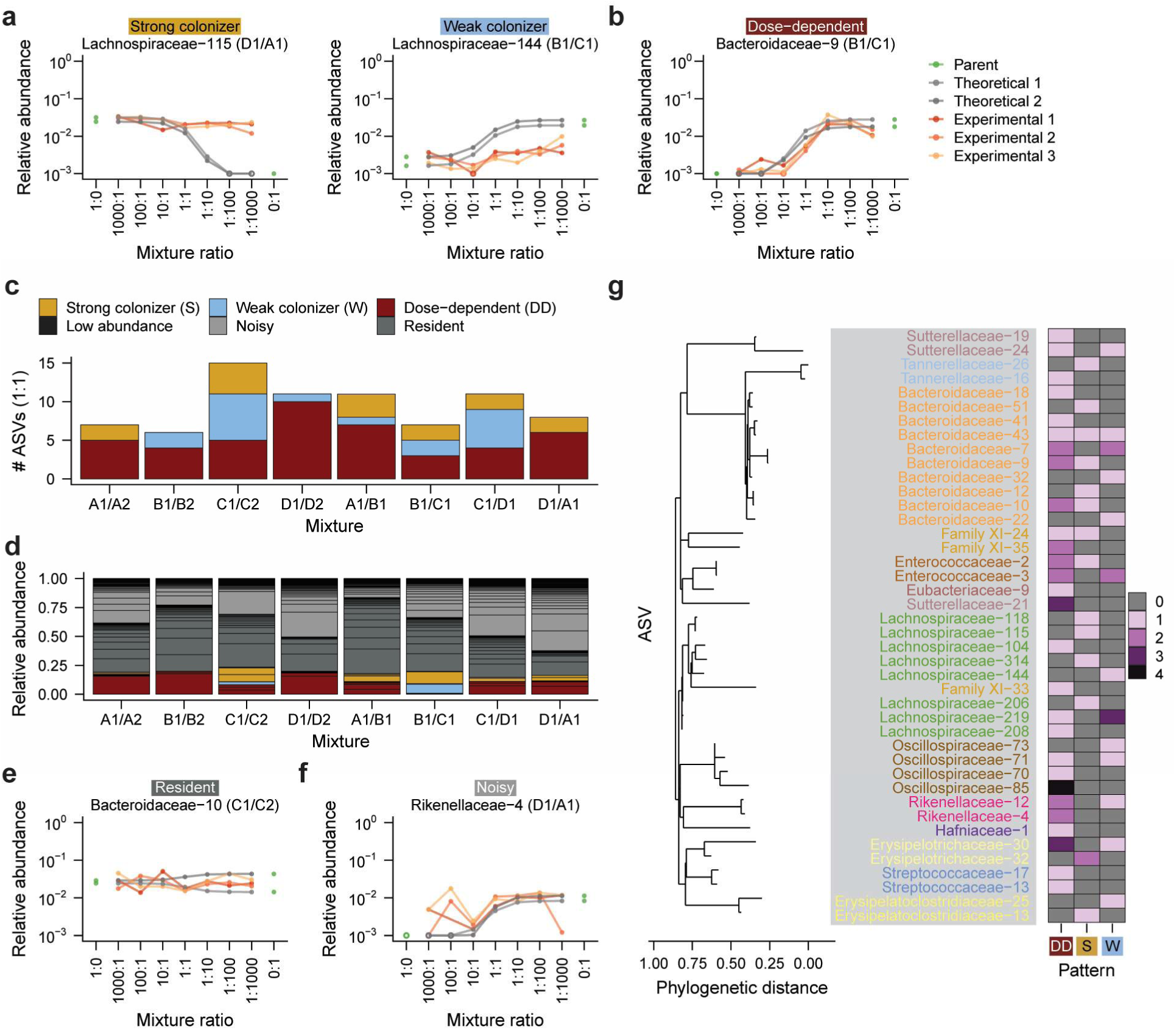
Colonization behaviors of individual ASVs influence community composition. a) Representative examples of strong and weak colonizers. Each panel shows an individual ASV from one mixture. Gray lines represent both replicates of predicted neutral relative abundances for each ASV, and orange lines represent three replicates of experimental relative abundances. b) Dose-dependent colonizers exhibit large changes in relative abundance across mixture ratios. Lines are colored as in (**a**). c) The number of ASVs that displayed a strong, weak, or dose-dependent colonization pattern in each set of mixtures. ASVs were counted as present if they were detected in the 1:1 mixture at the fifth passage. d) Total relative abundance of each type of colonizer in each set of mixtures. Only the 1:1 mixture is shown. Black lines indicate separate ASVs. Relative abundances are averaged across the three inoculation replicates. e) A representative example of a resident ASV that was present at similar relative abundances in both parent communities. Lines are colored as in (**a**). f) A representative example of a noisy ASV whose relative abundance showed large, non-monotonic fluctuations between adjacent mixture ratios. Lines are colored as in (**a**). g) Colonization behavior is not associated with ASV phylogeny. The heatmap shows the number of times each ASV exhibited a dose-dependent (DD), strong (S), or weak (W) colonization pattern in any mixture.

In contrast, many of the remaining ASVs exhibited dose-dependent colonization, with relative abundances that varied systematically across mixture ratios (*n*=43/410, 10.5%, **Fig. 2b, Extended Data Fig. 6, 7c**). Dose-dependent colonizers were present in every community mixture (**Fig. 2c**) and accounted for up to 20% of co-culture relative abundance at the 1:1 mixture ratio (**Fig. 2d**). Most of the remaining abundance was composed of resident ASVs, which were present at approximately the same relative abundance in both parent communities and across all mixture ratios (*n*=175/410, 42.7%, **Fig. 2e, Extended Data Fig. 6, 8a**), or by noisy ASVs whose relative abundance changed non-monotonically across mixture ratios (*n*=160/410, 39.0%, **Fig. 2f, Extended Data Fig. 6, 8b**). The prevalence of dose dependence among these varied colonization behaviors demonstrates that propagule size can influence the colonization of a sizeable fraction of species in each community during coalescence, even when communities show substantial deviations from neutral composition (**Fig. 1e**, **Fig. 2c,d**). Intriguingly, most ASVs exhibited distinct colonization patterns across co-cultures and relative to closely related taxa (**Fig. 2g, Methods**), suggesting that these patterns were influenced more strongly by community context than by species identity or phylogeny. Taken together, these behaviors reveal that species in the same co-cultures can exhibit varying levels of dependence on initial conditions.

### Transient dose dependence arises from high niche overlap in a consumer-resource model

We next sought to determine how the combination of neutral and non-neutral forces in each mixture leads to dose-dependent colonization patterns. In the D1/D2 co-cultures, we identified three dose-dependent *Lactococcus* and *Enterococcus* colonizers from the D1 parent community. The relative abundances of these ASVs were approximately anticorrelated with those of another dose-dependent *Lactococcus* from the D2 community (**Fig. 3a**). This observation led us to hypothesize that dose dependence could arise from competition between phylogenetically related taxa that occupy overlapping niches.

**Figure 3:**
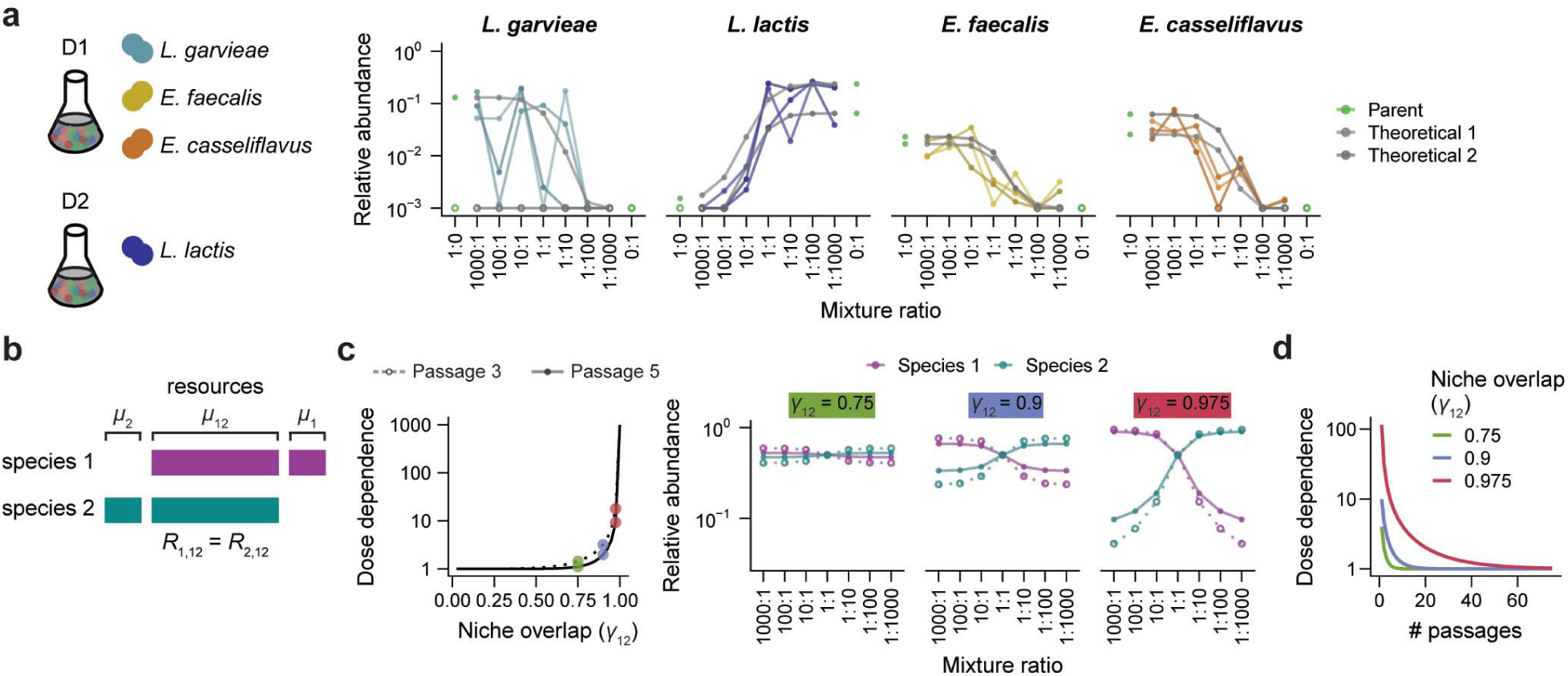
A consumer-resource model predicts transient dose dependence. a) ASVs from the *Enterococcus* and *Lactococcus* genera showed dose dependence during community coalescence. *Enterococcus faecalis*, *Enterococcus casseliflavus*, and *Lactococcus lactis* were isolated from the D1 community, and *Lactococcus lactis* was isolated from the D2 community. Gray lines represent both replicates of predicted neutral relative abundances for each ASV after five passages, and colored lines represent three replicates of experimental relative abundances after five passages. b) Schematic of a simple consumer-resource model in which two species compete neutrally (equal consumption rates *R*_1,12_ = *R*_2,12_ = 1) for resource μ_12_, and resources μ_1_ and μ_2_ are unique to species 1 and 2, respectively (**Methods**). c) The dose dependence of both species in the consumer-resource model increases as the niche overlap of species 1 with species 2 (γ_12_) increases. Dose dependence also decreases from passage 3 (dotted line) to passage 5 (solid line). Colored points highlight three values of γ_12_ for which the relative abundances over mixture ratios of each species are shown at right. Dose dependence is defined as the ratio of relative abundances after passaging starting from a 1000:1 versus 1:1000 mixture ratio. d) Dose dependence declines over passages in the two-species consumer resource model, even when niche overlap (γ_12_) is high.

We used a standard consumer-resource model^19^ to investigate the conditions under which competition for shared resources could give rise to neutral-like colonization (**Methods**). We first studied a system in which two species compete for a set of three common resources: μ_1_, μ_2_, and μ_12_ (**Fig. 3b**). Two of these resources, μ_1_ and μ_2_, are consumed exclusively by species 1 and 2, respectively, and the third resource, μ_12_, is consumed by both species at equal rates (*R*_1,12_ = *R*_2,12_). We investigated how the niche overlap of species 1 with species 2, defined as 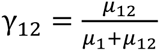, would affect the colonization behavior of these two species. We varied γ_12_ from 0.025 to 1 while maintaining the same amount of unique resources for each species (μ_1_ = μ_2_). We initialized the abundances of the two species at ratios from 1000:1 to 1:1000, as in our community coalescence experiments, and we simulated the model to test how species abundances would change over five passages.

In this system, the dose dependence of both species increased with the amount of overlap between their niches (**Figure 3c**). For example, when niche overlap was low (γ_12_ = 0.75), the two species rapidly reached consistent relative abundances across mixture ratios (**Figure 3c**). By contrast, when niche overlap was high (γ_12_ = 0.975), the relative abundance of both species remained dose dependent, varying by ∼20-fold between the 1000:1 and 1:1000 mixture ratios after three passages (**Figure 3c**), consistent with the magnitude of dose dependence that we often observed during community coalescence (**Fig. 2b**, **Fig. 3a**). Similar levels of dose dependence were present even when the consumption rate of the shared resource varied by ∼10% relative to that of the unique resource (**Extended Data Fig. 9a**). These results show that neutral-like colonization can arise when the growth of two species is dominated by their consumption of shared resources.

Although these species showed dose dependence when niche overlap was high, the magnitude of this dose dependence declined over time (**Fig. 3c,d**). For example, when γ_12_ = 0.975, the dose dependence of the two species declined from ∼20-fold to ∼10-fold from passage three to passage five (**Fig. 3c**). This declining dose dependence was consistent with decreases in dose dependence over passages that we observed experimentally for some ASVs during community coalescence (**Extended Data Fig. 9b**). Taken together, our model suggests that high niche overlap can slow a community’s convergence to equilibrium, creating transient, neutral-like colonization dynamics.

### Strain isolates show less dose dependence in pairwise mixtures than during community coalescence

Based on the predictions of our consumer-resource model, we hypothesized that the *Lactococcus* and *Enterococcus* species that showed dose-dependent colonization in the D1/D2 co-cultures had high levels of niche overlap that would lead them to show neutral-like colonization in pairwise mixtures as well. To test this prediction, we isolated representatives of these ASVs (**Methods**). We performed Sanger sequencing of the 16S rRNA gene and whole-genome sequencing (**Methods**) to identify the strains from the D1 community as *Lactococcus garvieae*, *Enterococcus faecalis*, and *Enterococcus casseliflavus*, and the strain from the D2 community as *Lactococcus lactis* (**Fig. 4a**).

**Figure 4:**
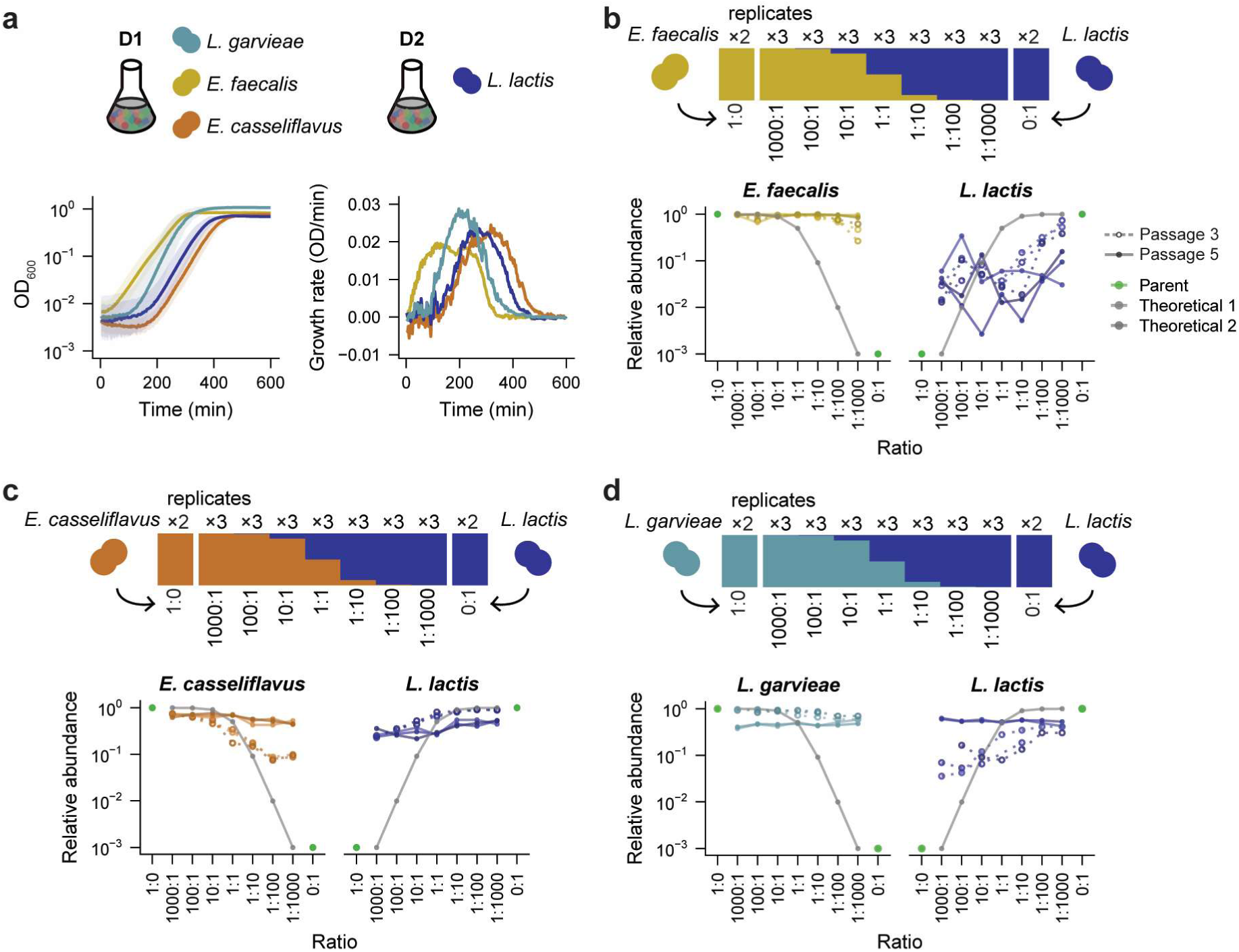
Transient dose-dependent colonization in pairwise strain mixtures. a) Growth of the *Enterococcus* and *Lactococcus* strains in monoculture. Instataneous growth rates (bottom right) were calculated from blank-subtracted growth curves (bottom left) after smoothing over a 5-min window (**Methods**). b-d) Species relative abundances in co-cultures of (b) *E. faecalis* and *L. lactis*, (c) *E. casseliflavus* and *L. lactis*, and (d) *L. garvieae* and *L. lactis*. Lines are colored as in Fig. 3a. Dose dependence decreased substantially from passage three (dotted lines) to passage five (solid lines).

These strains exhibited similar maximum growth rates in monoculture, although small differences in growth dynamics suggested the potential for distinct profiles of resource utilization (**Fig. 4a**).

We performed a set of pairwise mixtures of *L. lactis* (from community D2) with the strains isolated from community D1 at ratios from 1000:1 to 1:1000 (**Fig. 4b-d**). Most strains exhibited some degree of dose-dependent colonization after three passages, suggesting that their niches overlapped to some degree (**Fig. 4b-d**). However, after five passages, each strain reached a constant relative abundance across mixture ratios (**Fig. 4b-d**). This lack of dose dependence contrasted with their behavior in the D1/D2 co-cultures, in which dose dependence was maintained even after five passages (**Fig. 3a**). This discrepancy suggests that pairwise strain mixtures are unlikely to exhibit the same magnitude and duration of neutral-like colonization that we observed during coalescence of more complex communities.

### Dose dependence is enhanced in mixtures of more diverse communities

We used our consumer-resource model to investigate how colonization dynamics would be affected by the addition of a third species with overlapping niches. We introduced a third species into the model that consumes a unique resource initialized each passage at the same level as the other resources (i.e., μ_3_ = μ_2_, **Fig. 5a**). Species 3 also competes with species 1 for a fraction *β* of resource μ_1_, which it consumes at relative rate *r* compared to species 1. We varied *β* while keeping *r* constant (**Fig. 5b**) and vice versa (**Extended Data Fig. 9c**), initializing the abundances of species 1 and 2 at a range of ratios from 1000:1 to 1:1000 as before, and we set the abundance of species 3 equal to that of species 2 as if they were members of the same community. We simulated the model to investigate how species abundances changed over five passages.

**Figure 5:**
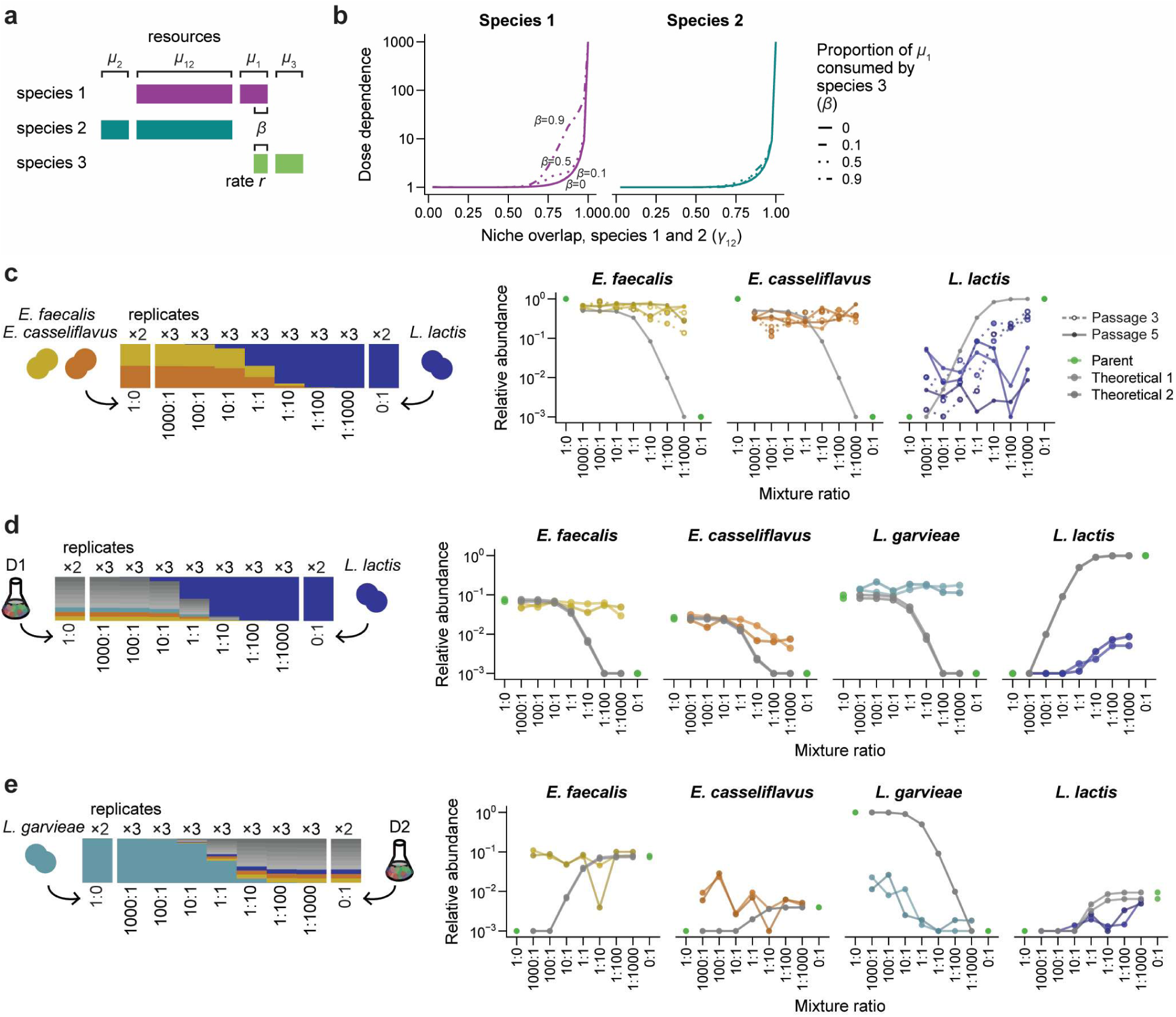
Dose dependence increases in mixtures with more diverse communities. a) Modification to the consumer-resource model in Fig. 3a in which a third species competes with species 1 for a fraction *β* of resource μ_1_ with relative consumption rate *r* compared to species 1. b) The dose dependence of species 1 after five passages increases with increasing niche overlap between species 1 and 3 (β) at constant *r* = 10, while the dose dependence of species 2 is largely unaffected. Dose dependence is defined as in Fig. 3c. c-e) Species relative abundances in co-cultures of (c) *E. faecalis, E. casseliflavus*, and *L. lactis*, (d) the D1 community and *L. lactis*, and (e) *L. garvieae* and the D2 community. Lines are colored as in Fig. 3a. Dotted lines with open points show data after three passages, and solid lines with closed points show passage five.

In this three-species model, species 1 showed higher dose dependence as its niche overlap with species 3 increased (**Fig. 5b**). When species 3 had minimal niche overlap with species 1 (*β* = 0.1, **Fig. 5b**) or was a weak competitor (*r* ≪ 1, **Extended Data Fig. 9c**), species 1 showed similar levels of dose dependence as when species 3 was absent (**Fig. 3c**), as expected. However, when niche overlap was high (*β* > 0.5) and species 3 was a strong competitor (*r* ≫ 1), species 3 effectively outcompeted species 1 for the use of their shared resources (**Fig. 5b, Extended Data Fig. 9c**). Under these conditions, the dose dependence of species 1 increased substantially due to its increased reliance on the resource shared with species 2 (**Fig. 5b**, **Extended Data Fig. 9c**). At the same time, the dose dependence of species 2 was essentially unaffected by the amount of niche overlap between species 1 and 3, showing that the addition of new species does not necessarily alter the colonization behavior of all community members (**Fig. 5b, Extended Data Fig. 9c**). Together, these results predict that neutral-like colonization dynamics should be longer-lasting and more pronounced in diverse communities, in which competition with multiple species limits the unique resources available for any species.

To test this prediction, we introduced the *L. lactis* strain into an equal mixture of *E. faecalis* and *E. casseliflavus* at ratios from 1000:1 to 1:1000. Consistent with our prediction, after three passages, the degree of dose dependence exhibited by *L. lactis* increased to ∼200-fold (**Fig. 5c**) from ∼60-fold in pairwise co-cultures (**Fig. 4b,c**), while neither *E. faecalis* nor *E. casselifavus* exhibited dose-dependent colonization (**Fig. 5c**). After five passages, none of the three strains exhibited dose dependence (**Fig. 5c**). These results support our model’s prediction that the addition of a new species can create more pronounced neutral-like colonization dynamics, even if dose dependence still declines more rapidly than during coalescence of complex communities.

To test the prediction that introducing strains into a more diverse community would increase the magnitude and duration of dose dependence, we performed a set of strain-community mixtures in which we mixed each of the two *Enterococcus* and *Lactococcus* strains into the D1 or D2 communities at ratios from 1000:1 to 1:1000 (**Fig. 5d,e, Extended Data Fig. 10**). In striking contrast to the pairwise and three-strain mixtures, a subset of strains in each strain-community mixture showed dose dependence after five passages (**Fig. 5d,e, Extended Data Fig. 10**). For instance, *E. casseliflavus* and *L. lactis* were no longer dose dependent after the fifth passage in pairwise co-cultures (**Fig. 4c**). However, when *L. lactis* was introduced into the D1 community, both strains exhibited dose dependence after five passages, varying by ∼10-fold in relative abundance across all mixture ratios (**Fig. 5d**). Similarly, when *L. garvieae* was introduced into the D2 community, it showed complementary patterns of dose dependence with *L. lactis*, even though *E. casseliflavus* remained at constant relative abundance across mixture ratios (**Fig. 5e, Extended Data Fig. 10c**). Together, these results demonstrate that dose dependence can increase with community diversity. These findings are consistent with the model prediction that the presence of additional species increases niche overlap, prolonging the effect of initial propagule size and creating neutral-like colonization dynamics.

## Discussion

In this study, we quantified the effect of propagule size on hundreds of simultaneous colonization events during community coalescence, and we compared the resulting co-cultures to neutral ecological predictions. Community composition deviated systematically from neutral predictions, but initial population size had a strong effect on colonization for a subset of species in each mixture, even after ∼40 generations of competition. Using a consumer-resource model, we showed that this dose-dependent colonization can arise from non-neutral competition for shared resources. These neutral-like dynamics frequently emerge in diverse communities due to high niche overlap among community members, even in the absence of more complex phenomena like resource fluctuations^49,50^, spatial structure^51,52^, and environmental modification^7,14^.

Our model provides an explanation for how neutral-like colonization can occur in a community dominated by non-neutral competition. When considering either neutral theory^18,53^ or resource competition alone^49,54^, all species in a community are expected to show the same colonization dynamics. However, we observed both neutral-like and non-neutral colonization dynamics in each community mixture. Our model explains this co-occurrence by showing that neutral-like behavior is neither an intrinsic property of a species, nor of the community as a whole. Instead, neutral-like dynamics are emergent phenomena^55–58^ that arise from non-neutral competitive interactions among subsets of community members.

Another feature of our model is that neutral-like colonization dynamics are transient: communities eventually converge to a stable equilibrium, as in classical consumer-resource models^19^. However, both our model and experiments show that neutral-like dynamics can persist for ∼40 generations. The timescales of these transient states^59–61^ may be long enough for introduced species to affect community composition and function^60,62,63^, induce shifts between alternative states^50,64^, and alter the outcomes of subsequent species introductions^14,65–68^—even if the initial species introductions are ultimately unsuccessful.

Our findings have several implications for the design of ecological interventions like microbiome therapeutics^35^ and agricultural inoculants^69^. We showed that introduced species often have higher levels of dose dependence in diverse communities, meaning that experiments with small, model communities may systematically underestimate the effects of propagule size in more diverse, natural contexts. Our model also explains why the same species can show varying levels of dose dependence in different community contexts, which may explain why previous studies have come to contradictory conclusions about how propagule size affects colonization success^5,20,26,30,33,34,37^.

This work identifies niche overlap as the main reason propagule size has stronger impacts on colonization in more diverse communities. While niche overlap is challenging to measure, it is often higher among closely related species, which tend to consume more similar resources^70–72^. We might therefore expect higher levels of dose dependence when introduced species are taxonomically related to existing residents, as our *Lactococcus* and *Enterococcus* example illustrates. However, more precise methods to quantify resource utilization, such as metabolomics^22,71,73,74^, may be necessary to accurately predict niche overlap in microbial communities. Future studies that combine controlled colonization experiments with metabolomics of microbial isolates will improve our ability to predict how neutral forces like propagule size impact colonization.

## Materials and Methods

### Data availability

Sequencing data will be made publicly available from the NCBI SRA upon manuscript publication.

## Code availability

The computer code that performs the analysis is available at https://github.com/DoranG1/dose-dependent-colonization.

### Collection of stool samples from human subjects

Stool samples were collected as a part of a study^75^ approved by the Stanford University Institutional Review Board under Protocol 54715. Informed written consent was obtained from all participants. Samples were provided by healthy, non-pregnant adults living with at least one roommate or housemate. At least two participants were recruited to provide samples from each household, one of whom took a five-day course of the antibiotic ciprofloxacin during sampling. Subjects collected stool samples in sterile plastic tubes and froze them immediately at -20°C inside multiple layers of insulating protection. Samples were transported to the laboratory on dry ice within six weeks of collection and were stored at -80°C. For this study, we derived *in vitro* communities from the last pre-antibiotic stool sample collected by eight study subjects from four household pairs. These eight subjects lived in the San Francisco Bay Area and ranged from 25-30 years old.

### Derivation of top-down *in vitro* communities

To obtain stool fragments for inoculation of *in vitro* communities, we used a Basque Engineering CXT 353 Benchtop Frozen Aliquotter to drill cores from the frozen stool samples of eight study subjects: A1, A2, B1, B2, C1, C2, D1, and D2 (**Table S1**). Cores were kept frozen by immersing the drill area in liquid nitrogen and were stored at -80 °C after drilling.

We generated *in vitro* communities following methods established in our previous studies^46,47^. All steps of community inoculation and passaging were performed in an anaerobic chamber (Coy Instruments), using sterilized pipette tips. First, a 50-mg fragment of each stool core was resuspended in 500 µL of filter-sterilized PBS. Next, 20 µL of the stool resuspension were added to 180 µL of Brain Heart Infusion (BHI, BD Biosciences 237200) supplemented with 0.2 mg/mL L-tryptophan, 1 mg/mL L-arginine, 0.5 mg/mL L-cysteine, 5 µg/mL vitamin K, 1 µg/mL haemin, and 5 g/mL mucin (referred to here as mBHI+mucin) in a flat-bottom 96-well plate (Greiner Bio-One 655161). Each sample was inoculated as three replicates in separate wells. All media were autoclaved before inoculation to ensure sterility.

After inoculation, communities were grown at 37 °C in the anaerobic chamber and passaged every 48 h by transferring 1 µL of saturated culture into a new 96-well plate containing 199 µL of fresh mBHI+mucin. We passaged each community 15 times and stored each plate of saturated cultures at -80 °C after passaging by mixing 100 µL of saturated culture with 100 µL of 50% filter-sterilized glycerol.

### Whole community mixtures

We thawed glycerol stocks of one replicate of each community from passage 15 and reinoculated these cultures in an anaerobic chamber (Coy Instruments) by adding 3 µL of glycerol stock to 197 µL of mBHI+mucin. These communities were grown at 37 °C for 48 h.

To create community mixtures, we first generated a dilution series of each of the unmixed parent communities. Fresh mBHI+mucin was added to each saturated culture to create communities with concentrations 1/10, 1/100, and 1/1000 of the parent community. We generated each dilution in triplicate from the single replicate of each parent community, resulting in 24 communities for each dilution factor from the eight parent communities, and 72 dilution communities in total across all three dilution factors.

To generate each pairwise mixture, we added 1 µL of each dilution community to 1 µL of another parent community and 198 µL of fresh mBHI+mucin. We mixed communities in eight combinations: A1/A2, B1/B2, C1/C2, D1/D2, A1/B1, B1/C1, C1/D1, and D1/A1. Four pairs of communities (A1/A2, B1/B2, C1/C2, D1/D2) were derived from subjects from the same household, while the other four mixtures (A1/B1, B1/C1, C1/D1, D1/A1) involved communities from different households. Each pair of communities was mixed at seven ratios: 1000:1, 100:1, 10:1, 1:1, 1:10, 1:100, and 1:1000. Each mixture was passaged in triplicate. This protocol resulted in 168 co-culture communities (8 mixtures × 7 ratios × 3 replicates), which were passaged in two flat-bottom 96 well plates. We also inoculated 2 µL of each undiluted parent community in 198 µL of mBHI+mucin in duplicate, comprising 16 unmixed control parent communities. The four corners of each 96-well plate were filled with 200 µL of mBHI+mucin as blank controls to monitor contamination.

Both 96-well plates containing mixture co-cultures and unmixed control communities were grown at 37 °C and passaged every 48 h five times by adding 1 µL of saturated culture to 199 µL of mBHI+mucin. Saturated cultures from each passage were stored at -80 °C.

### 16S rRNA gene sequencing

DNA was extracted from 50 µL of each community from each mixture using the Qiagen DNeasy Ultraclean 96 Microbial Kit (Qiagen 10196-4) according to the manufacturer’s instructions. Extracted DNA was stored at -20 °C.

We performed 16S rRNA gene amplicon sequencing targeted to the V4 region. Extracted DNA was amplified using primers modified from the Earth Microbiome Project spanning 515F–806R, with gene-specific sequences GTGYCAGCMGCCGCGGTAA for the forward primer and GGACTACNVGGGTWTCTAAT for the reverse primer^76,77^. The 16S region was amplified using 27 cycles of PCR with an annealing temperature of 50 °C, and successful amplification of the 16S region was confirmed using gel electrophoresis. Sequencing primers were attached using 10 cycles of PCR with an annealing temperature of 54 °C. Excess primer was removed using a 0.8X Ampure bead cleanup. Libraries were pooled to contain equal volume from each sample and sequenced using an Illumina NovaSeq SP at the DNA Services Lab, Roy J. Carver Biotechnology Center, University of Illinois at Urbana-Champaign.

Raw reads were annotated and demultiplexed using UMI-tools^78^, and primer and adapter sequences were trimmed using cutadapt^79^. To remove the blank media controls, samples with <100 reads were excluded. DADA2^80^ was used to filter and truncate reads and to assign amplicon sequence variant (ASV) taxonomy based on the SILVA (release 138) reference database^81^, and to obtain a phylogeny of all ASVs.

Our final community coalescence dataset contained 91,504,441 reads with a median of 201,425 reads (standard deviation 69,902) per non-blank sample. Two samples in our dataset (A1 replicate 1 from passage 3, and D1/A1 100:1 replicate 1 from passage 5) received zero sequencing coverage, so these were excluded prior to subsequent analyses. 1715 unique ASVs were detected across all samples, and 198 of these were present at ≥10^-3^ relative abundance in at least one sample, which we estimated as the limit of detection. Although subjects from the same households often shared a higher proportion of gut microbial strains compared to subjects from different households^75^, we observed no differences in the proportion of ASVs shared in mixtures of communities from the same versus different households (**Extended Data Fig. 3**). We used the relative abundance of each ASV to calculate Jensen-Shannon divergence.

In addition to our community coalescence dataset, we obtained a second dataset representing the *L. garvieae*/*L. lactis* co-cultures after five passages and the D2/*L. lactis* co-cultures after five passages. This dataset contained 48,347,562 reads with a median of 683,158 reads (standard deviation 370,504) per non-blank sample. We also obtained a third dataset representing the *E. faecalis*/*L. lactis*, *E. casseliflavus*/*L. lactis*, three-strain, and all strain-community co-cultures after three and five passages. This third dataset contained 9,429,310 reads with a median of 12,927 reads (standard deviation 12,375) per non-blank sample. We excluded one replicate each from the *L. garvieae*/*L. lactis*, D1/*L. lactis, L. garvieae*/D2, and *E. casseliflavus*/D2 (passage 5) mixtures based on poor sequencing coverage at one or more mixture ratios in those replicates.

### Estimation of theoretical community compositions

To assess the effect of propagule size in our community mixtures, we computationally generated a set of theoretical community mixtures according to the predictions of ecological neutral theory^18^. The expected abundance of each ASV across mixture ratios was calculated based on a weighted average of the relative abundance in the parent communities: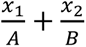 where *x*_1_ and *x*_2_represent the relative abundance of the ASV in parent community 1 and 2, respectively, and *A* and *B* represent the dilution factor of parent community 1 and 2, respectively, at the given mixture ratio. For example, the predicted abundance of an ASV in a 1:100 mixture of A1 and A2 in which the propagule size of A2 is 100 times greater than that of A1 is 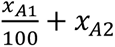. We normalized these weighted averages to calculate a theoretical relative abundance for each ASV in a mixture, and we ignored ASVs with predicted relative abundance <10^-3^, our sequencing limit of detection. We generated two theoretical communities for each pair of parent communities based on the two replicates of each parent community in our experiments. We generated separate sets of theoretical communities from passages 3 and 5 of the parent communities to account for changes to parent community composition that may have occurred during passaging.

### Categorization of ASVs by colonization behavior

We developed a set of summary statistics for each ASV based on its average change in relative abundance across mixture ratios and the deviation from predicted neutral relative abundances, which enabled us to systematically categorize each ASV into one of six distinct patterns: low abundance, noisy, resident, dose-dependent, strong colonizer, or weak colonizer (**Extended Data Fig. 5**).

For each ASV in each mixture, we calculated four measures. The average log_10_ fold change in relative abundance across mixture ratios (net dose difference) and average absolute value of log_10_ fold change in relative abundance across mixture ratios (magnitude dose difference) were used to quantify each ASV’s level of dose dependence. The average log_10_ fold difference between experimental and theoretical relative abundances across mixture ratios (net theoretical difference) and average absolute value of log_10_ fold difference between experimental and theoretical relative abundances across mixture ratios (magnitude theoretical difference) were used to quantify each ASV’s deviation from neutral predictions and distinguish between resident, strong colonizing, and weak colonizing ASVs.

Net dose difference was calculated as 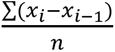, where *x_i_* represents the log_10_(relative abundance) of an ASV at a single mixture ratio (e.g., 100:1), *x*_i-1_ represents the log_10_(relative abundance) at the preceding mixture ratio (e.g., 1000:1), and *n* = 7 represents the total number of mixture ratios. Magnitude dose difference was calculated as 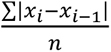. Both net dose difference and magnitude dose difference were first calculated individually for each replicate of the ASV and then averaged across replicates.

Net theoretical difference was calculated as 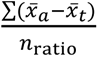, where 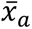 and 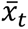 represent the average experimental and theoretical, respectively, log_10_(relative abundance) across replicates at a single mixture ratio. Magnitude theoretical difference was calculated as 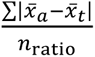. For ASVs that were not detected in the co-cultures at some mixture ratios and/or parent communities, we set the relative abundance to 10^-3^.

To categorize ASVs, we used five binary decisions based on our summary statistics and other measures (**Extended Data Fig. 5**). First, ASVs were classified as “low abundance” if their relative abundance was <10^-2.6^, a threshold slightly above our limit of detection, in both parent communities. Next, ASVs were classified as “noisy” if the ratio of magnitude dose difference to the absolute value of net dose difference was >2.25, indicating large, non-monotonic changes in relative abundance over mixture ratios. To prevent ASVs with extremely low change over mixture ratios (absolute value net dose difference <0.1) from artificially inflating the number of noisy ASVs, we replaced these low values with 0.1 as a pseudocount.

If ASVs were not classified as low abundance or noisy, they were classified as “dose-dependent colonizers” if the absolute value of the net dose difference was >0.09, meaning that they exhibited large monotonic changes in relative abundance over mixture ratios. Of the remaining ASVs, if the magnitude theoretical difference was >0.3 and the absolute value of the net theoretical difference was ≥0.2, indicating a strong deviation from neutral predictions, they were classified as “strong colonizers” or “weak colonizers” based on whether the sign of net theoretical difference was positive or negative, respectively. All other ASVs, which showed little dose dependence or deviation from neutral predictions, were classified as “resident”. All numeric cutoffs between categories were chosen to minimize the number of ASVs classified as low abundance, noisy, resident, or low abundance, while ensuring that dose-dependent, strong, and weak colonizers could be readily distinguished by inspection.

### Consumer-resource model

We implemented a standard consumer-resource model^19^:

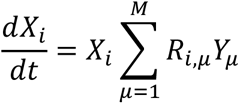

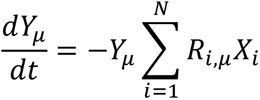

where *X*_i_ denotes the absolute abundance of species *i*, *Y*_µ_ the amount of resource μ, and *R*_i,µ_ the consumption rate of resource μ by species *i*. Species abundances were simulated in Matlab (Mathworks, Inc.) using the ode45 solver until all resources were depleted. After each such passage, the abundances were diluted 1:200, and resource concentrations were refreshed. We initialized the abundances of the two species at a range of ratios from 1000:1 to 1:1000, as in our community coalescence experiments, and we simulated five passages for each community. We defined dose dependence as the ratio of a species’ relative abundances at starting ratios of 1000:1 and 1:1000.

We first investigated a simple system in which two species, denoted 1 and 2, compete to consume use a set of three common resources: μ_1_, μ_2_, and μ_12_ (**Fig. 3b**). Two of these resources, μ_1_ and μ_2_, are consumed exclusively by species 1 and 2 respectively, and the third resource, μ_12_, is consumed by both species at equal rates (*R*_1,12_ = *R*_2,12_ = 1). To investigate how the niche overlap of species 1 with species 2, defined as γ_12_ = 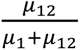, would affect the colonization behavior of these two species (**Fig. 3c**), we varied γ_12_ from 0.025 to 1 while maintaining the same amount of unique resources for each species μ_1_ = μ_2_ = 1. To test how varying the rates of shared resource consumption affects colonization behavior, we varied *R*_1,12_ from 0.9 to 1.1 while maintaining a constant *R*_2,12_ = 1 (**Extended Data Fig. 8a**).

We investigated how dose dependence would be affected by the addition of a third species. We introduced a third species into the model that consumes a unique resource, μ_3_ = μ_2_ = 1. Species 3 also competes with species 1 for a fraction *β* of resource μ_1_, which it consumes at relative rate 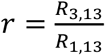 compared to species 1. To investigate how the presence of this new species affects colonization, we varied *β* from 0 to 0.9 while maintaining a constant *r* = 10 (**Fig. 5b**). As before, we initialized the abundances of species 1 and 2 at a range of ratios from 1000:1 to 1:1000, and we set the abundance of species 3 equal to that of species 2. To investigate how the consumption rate of this shared resource affects colonization behavior, we also tested the effect of varying *r* from 10^-2^ to 10^2^ while maintaining a constant *β* = 0.5 (**Extended Data Fig. 8d**).

### Strain isolation

*Lactococcus garvieae*, *Lactococcus lactis*, *Enterococcus faecalis*, and *Enterococcus casseliflavus* strains representing dose-dependent ASVs were isolated by sorting single cells from communities D1 and D2 into mBHI-filled wells of a 96-well plate using a fluorescence-activated cell sorter (FACS)^82^. To do so, communities were transported from the anaerobic chamber to the FACS facility in an anaerobic container (BD Biosciences) to ensure minimal oxygen exposure. We confirmed the purity of each strain isolate by streaking multiple times on BHI agar plates supplemented with 5% horse blood and submitted individual colonies for Sanger and whole-genome sequencing. Colonies verified as pure were grown for 48 h in 200 µL of mBHI+mucin in a 96-well plate, and 75 µL of each saturated culture were mixed with 75 µL of 50% glycerol and stocked in plastic crimp vials (ThermoFisher C4011-11) for future use.

To confirm isolate purity, we performed Sanger sequencing targeted to the V4 region of the 16S rRNA gene. We amplified the 16S gene with 35 cycles of PCR with an annealing temperature of 55 °C using the previously described primers. The resulting PCR products were sent to Elim Biopharmaceuticals for sequencing.

We determined the species identity of strain isolates using whole-genome sequencing. DNA was extracted from isolate cultures using the DNeasy Ultraclean 96 Microbial Kit (Qiagen 10196-4) according to the manufacturer’s instructions. Libraries were prepared for whole-genome sequencing using the Nextera DNA Flex Library Prep Kit (Illumina 20018705) according to the manufacturer’s instructions. The DNA concentration of each library was quantified using the Quant-iT High-Sensitivity dsDNA Assay Kit (Invitrogen Q33130) and a Synergy HT Microplate Reader (Biotek Instruments), and libraries were pooled at equal DNA input. Pooled libraries were sequenced at the DNA Services Lab, Roy J. Carver Biotechnology Center, University of Illinois at Urbana-Champaign.

### Co-cultures of strain isolates and strain-community mixtures

We performed a set of four mixtures of two or three strains (*L. garvieae*/*L. lactis*, *E. faecalis*/*L. lactis*, *E. casseliflavus*/*L. lactis*, and *E. faecalis*/*E. casseliflavus*/*L. lactis*) and five strain-community mixtures (D1/*L. lactis*, *L. garvieae*/D2, *E. faecalis*/D2, *E. casseliflavus*/D2, *E. faecalis*/*E. casseliflavus*/D2) using the glycerol stocks of our four strain isolates and of the D1 and D2 parent communities. We performed mixtures at ratios from 1000:1 to 1:1000 in triplicate as described above for whole community mixtures. For mixtures that included both *E. faecalis* and *E. casseliflavus*, 0.5 µL of each strain culture were used. Mixtures were passaged in mBHI+mucin for 48 h at 37 °C five times.

### Analysis of pairwise mixture composition using Sanger sequencing data

To analyze the composition of co-cultures from the *L. garvieae*/*L. lactis* mixture after three passages, we performed Sanger sequencing of each co-culture as described above and used the CASEU R package^83^ to quantify the relative abundance of both strains at each mixture ratio.

### Quantification of strain isolate growth

Growth curves were obtained using an Epoch 2 Microplate Spectrophotometer (Biotek Instruments). Strains were grown overnight and then diluted 1:10, 1:100, or 1:1000 into fresh BHI+mucin in 96-well plates. We measured the optical density at 600 nm (OD_600_) of 10 replicates of each strain over 48 h of anaerobic growth at 37 °C with continuous orbital shaking. Instantaneous growth rate was quantified as *d* ln(OD_600_)/*dt* using the gcplyr R package^84^.

## Author contributions

D.A.G., K.S.X., D.A.R., and K.C.H. designed the research; D.A.G., K.S.X., A.B.P., R.R.J., and L.R.F. performed the research. D.A.G., K.S.X., J.G.L., J.C.C.V., D.A.P., B.H.G., D.A.R., and K.C.H. analyzed the data; D.A.G., K.S.X., D.A.R., and K.C.H. wrote the paper; and all authors reviewed it before submission.

## Acknowledgements

We thank Mark Bitter, Tadashi Fukami, and members of the Huang, Relman, Good, and Petrov labs for helpful discussions. Sequencing support for this project was provided by the DNA Services Lab, Roy J. Carver Biotechnology Center, University of Illinois at Urbana-Champaign. This work was funded by Stanford Bio-X Undergraduate Summer Research Program fellowships (to D.A.G. and A.B.P.), Stanford Vice Provost for Undergraduate Education Small Grants (to D.A.G. and A.B.P.), a Carol Carmichael Summer Undergraduate Research Fellowship (to R.R.J.), NSF Awards IOS-2032985 and EF-2125383 (to K.C.H.), NIH Awards R01 AI147023 (to D.A.R. and K.C.H.) and RM1 GM135102 (to K.C.H.), the Thomas C. and Joan M. Merigan Endowment at Stanford University (to D.A.R.), and NIH/NIAID Award R21-AI168860 (to D.A.R.). J.G.L. is supported by the PRISM Baker Fellowship. K.S.X. has been supported by a James McDonnell Foundation Postdoctoral Fellowship in Understanding Dynamic and Multi-Scale Systems and a Jane Coffin Childs Memorial Fund Postdoctoral Fellowship. K.C.H., B.H.G., and D.A.P. are Chan Zuckerberg Biohub Investigators.

## Extended Data Figures

**Extended Data Figure 1:**
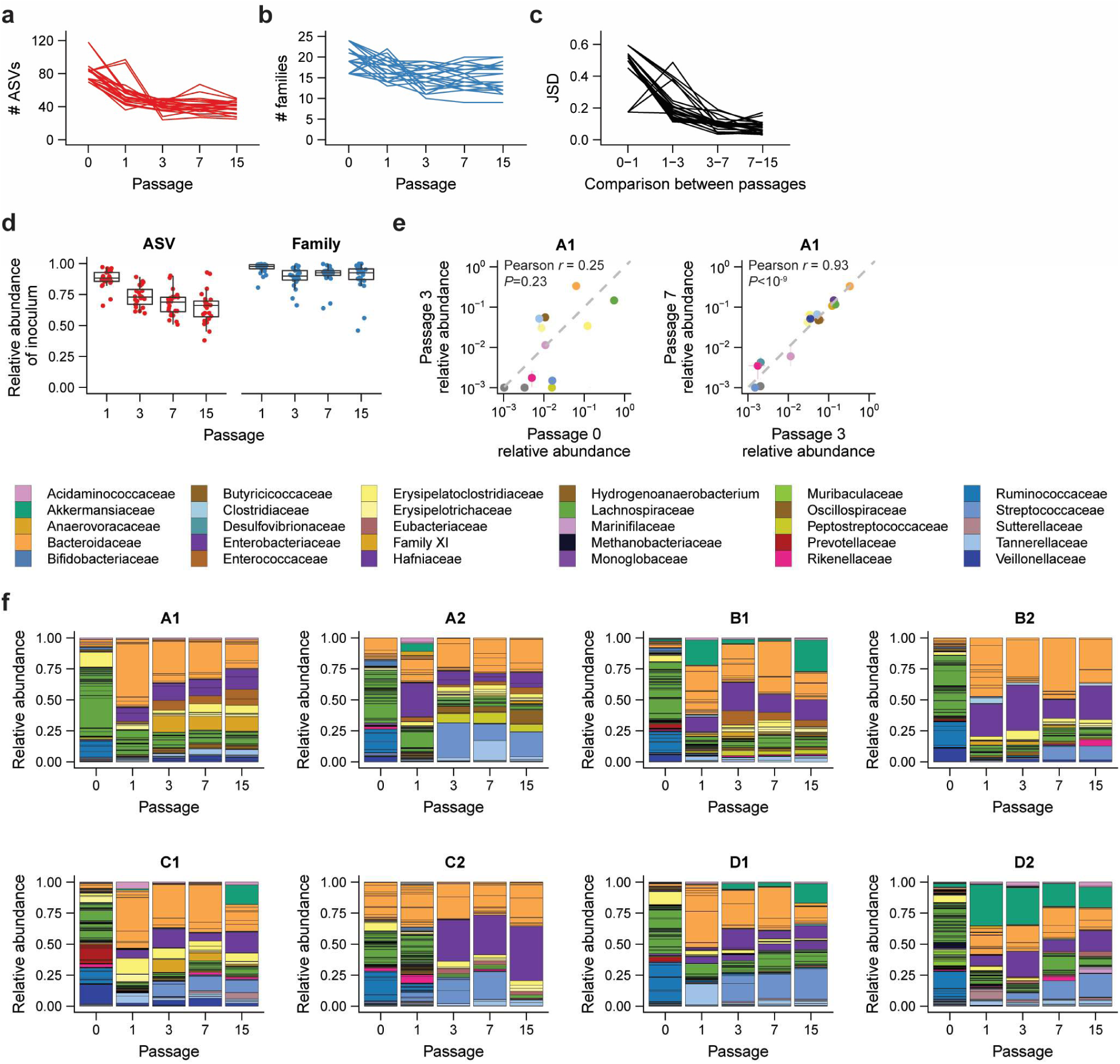
Diversity and composition of stool-derived communities. a) ASV richness of each community over passages. Only ASVs present in the community at relative abundance >10^-3^ were included in the richness calculation. b) Family-level richness of each community over passages. Families were counted as present if an ASV in the family was present in the community at relative abundance >10^-3^. c) Stability of each community over passages, as quantified by the Jensen-Shannon divergence (JSD, **Methods**) between consecutive passages. d) Proportion of the stool inoculum (passage 0) retained at each subsequent passage. Each point shows the total ASV or family relative abundance of the stool inoculum represented by taxa retained in communities at a given passage. Taxa were counted as retained if they were detected at a relative abundance >10^-3^. e) Correlation of family-level relative abundance across passages 0, 3, and 7 (Pearson’s tests, left: *n*=25, *ρ*=0.25, *P*=0.23; right: *n*=22, *ρ*=0.93, *P*<10^-9^). Points are colored by family. The dashed line indicates a perfect correlation in relative abundance between passages. f) Composition of one replicate of each community over passages. ASV relative abundances are separated with solid lines and colored by family. Only the most abundant families are shown in the key.

**Extended Data Figure 2:**
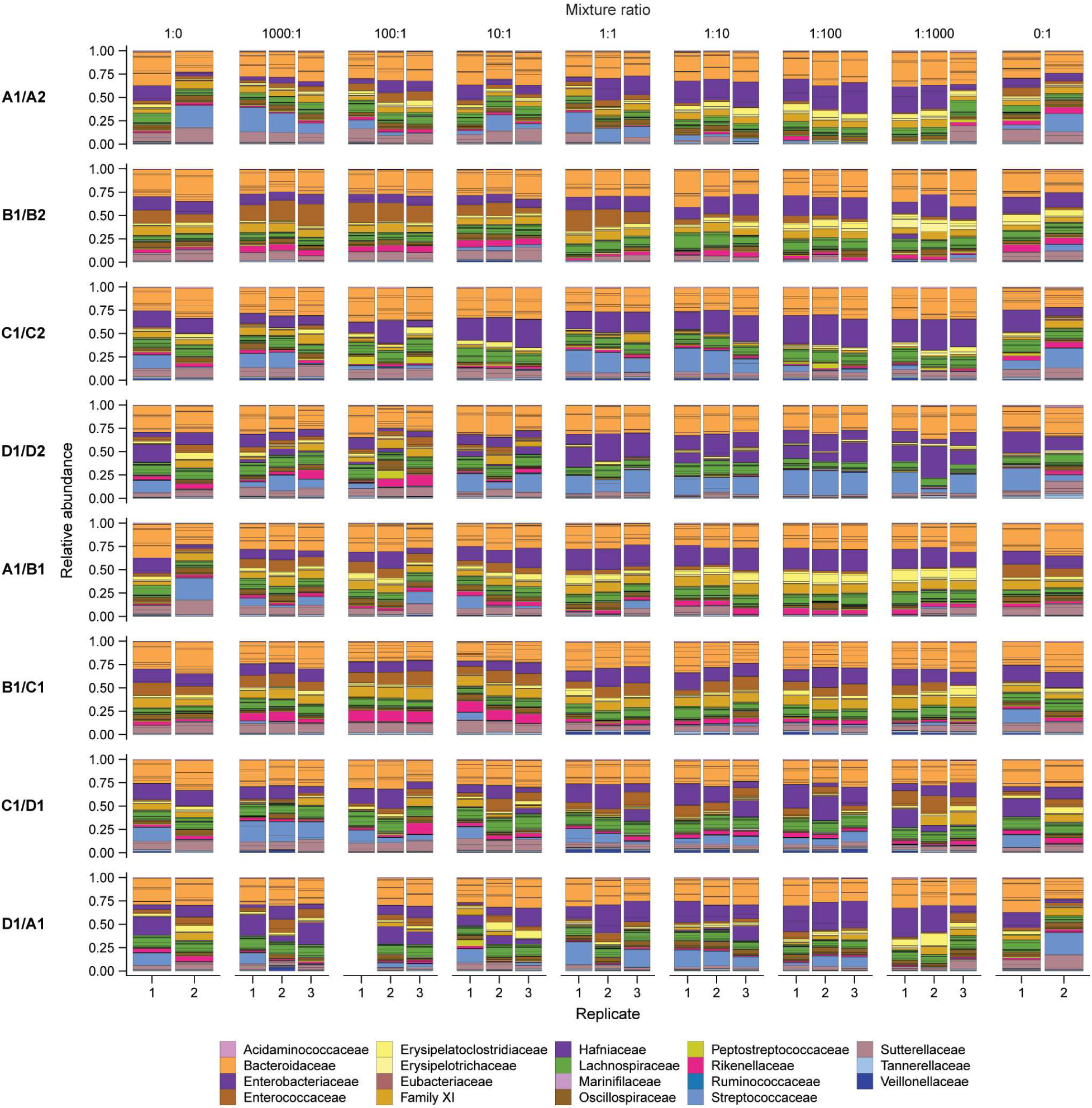
Composition of co-culture communities. ASV relative abundances are separated with solid lines and colored by bacterial family. Only the most abundant families are shown in the key. The first replicate of the 100:1 mixture ratio of the D1/A1 mixture had low sequencing depth and was removed from subsequent analyses.

**Extended Data Figure 3:**
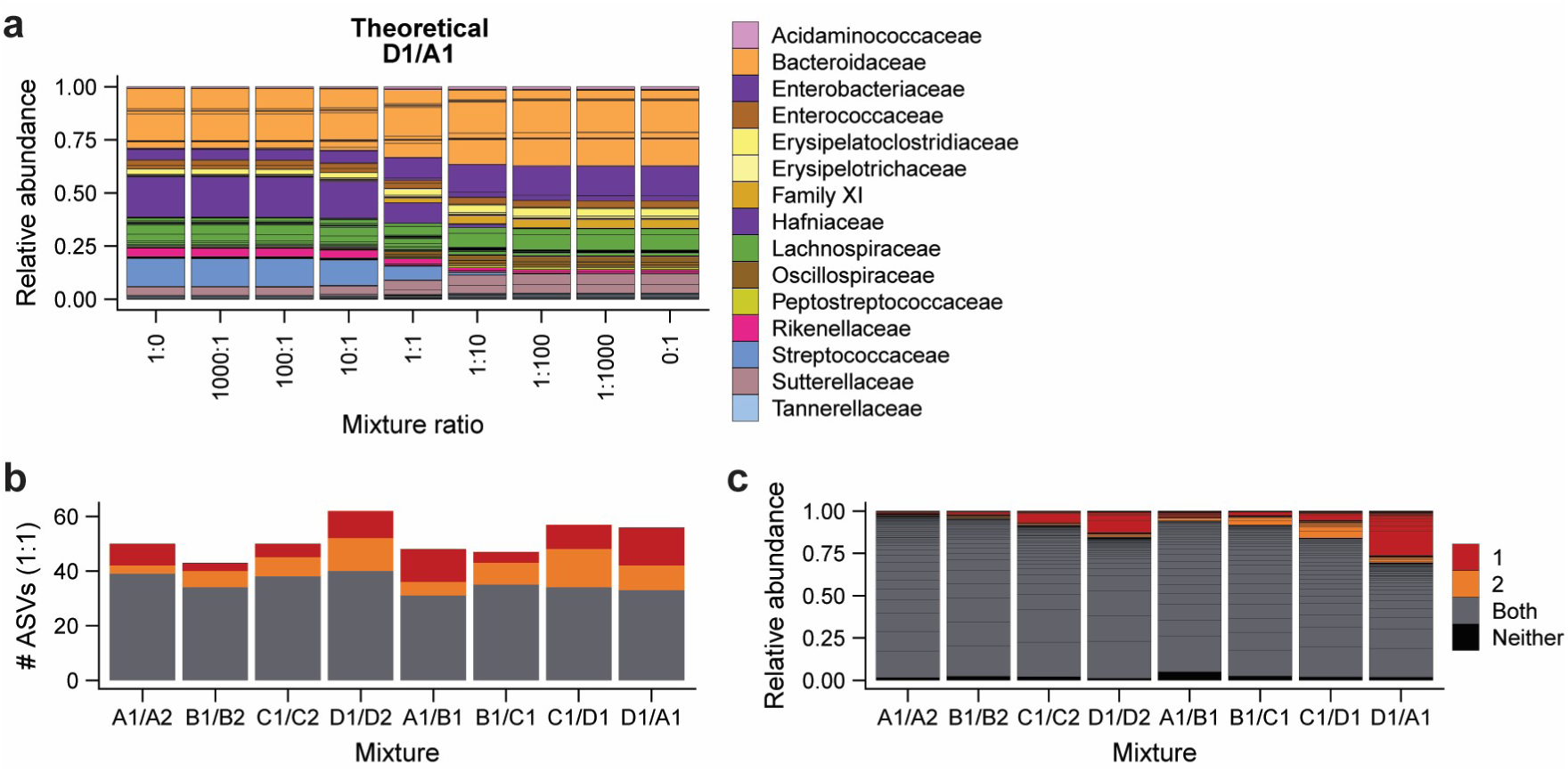
Theoretical community composition is dose dependent, even though many ASVs are shared between parent communities. a) The theoretical neutral community composition of an example mixture after five passages. The 1:0 and 0:1 ratios represent unmixed parent communities. ASV relative abundances are separated with solid lines and colored by family. b) Most ASVs in each co-culture were present in both parent communities. ASVs were counted as originating from one or both parent communities if they were present at relative abundance >10^3^. c) Relative abundances of ASVs at the 1:1 mixture ratio, colored by parent community of origin. Black lines indicate separate ASVs. Relative abundances were averaged across the three inoculation replicates.

**Extended Data Figure 4:**
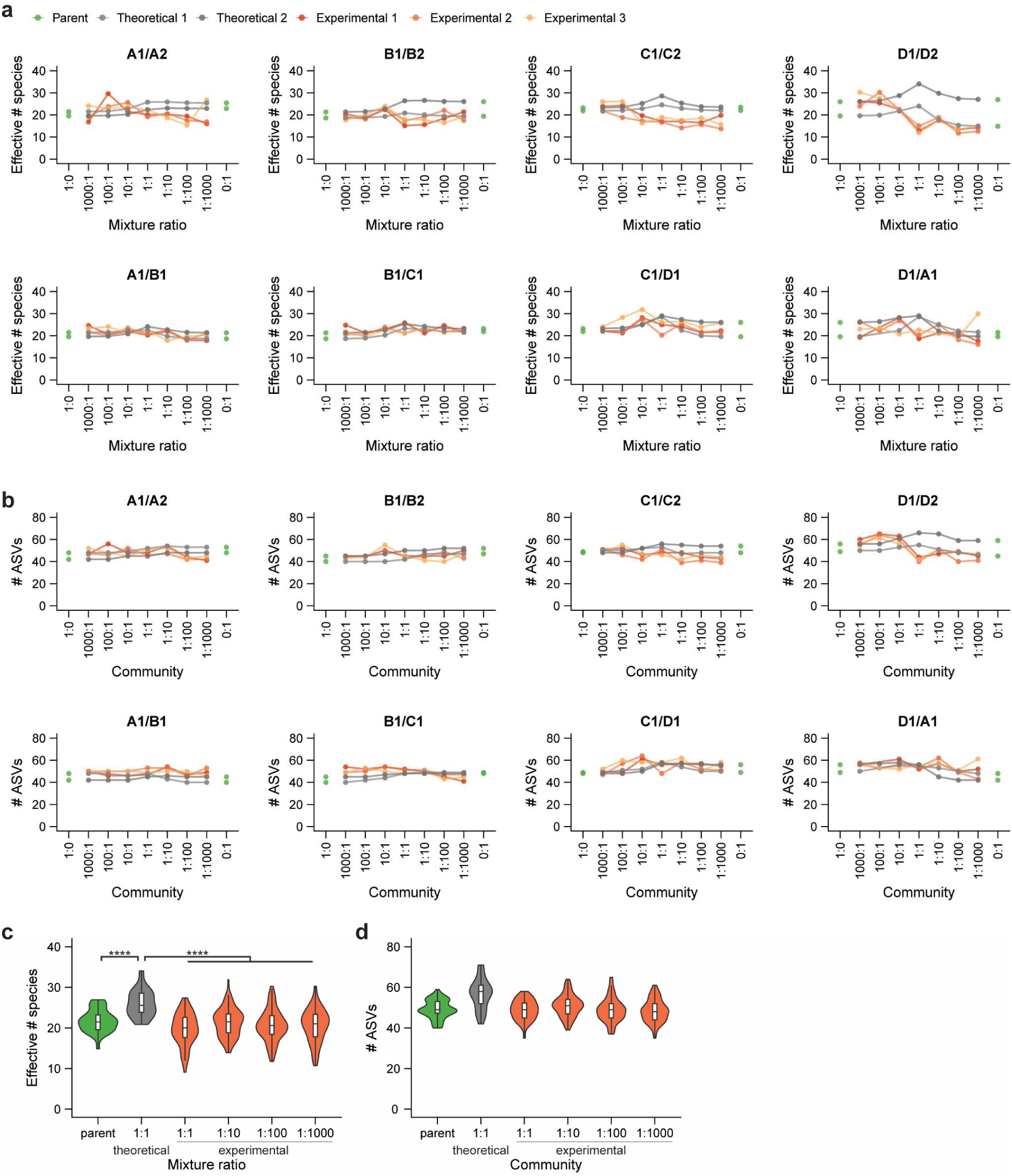
Diversity is limited in co-culture communities. a,b) Diversity of each co-culture community, quantified by (**a**) the effective number of species (*e*^H′^) calculated from the Shannon diversity index (*H*′), or (**b**) ASV richness, assuming a detection limit of 10^-3^ relative abundance. Lines are colored as in Fig. 1d. c) Distribution of diversity across parent and co-culture communities. The effective number of species at each mixture ratio more closely resembled the diversity of the parent communities than the higher-diversity theoretical prediction (Wilcoxon rank-sum one-sided tests, parent (*n*=32) vs. theoretical (*n*=16) co-cultures: *P*=6.1×10^-5^; theoretical vs. experimental 1:1 (*n*=24) co-cultures: *P*=6.6×10^-7^; theoretical vs. other (*n*=48 at each mixture ratio) co-cultures: *P*<10^-4^). d) Distribution of ASV richness across parent and co-culture communities.

**Extended Data Figure 5:**
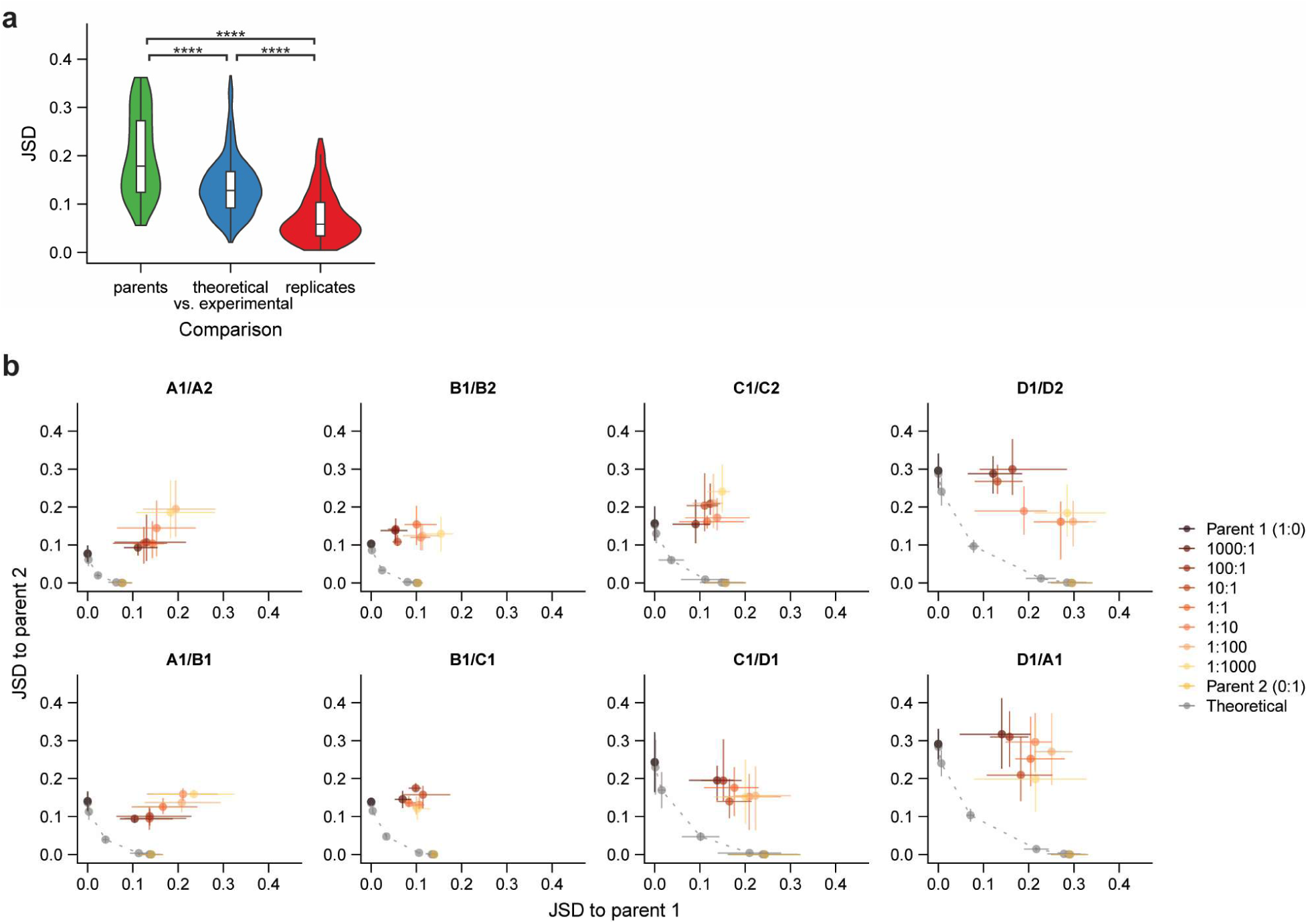
Composition of co-culture communities varies by mixture ratio but differs from neutral predictions. a) Comparisons of co-culture communities to theoretical communities. Violin plots represent the distribution of Jensen-Shannon divergence (JSD) between all pairs of parent communities (including those that were not mixed), theoretical and experimental mixtures at the 1:1 mixture ratio, and inoculation replicates of the 1:1 mixture ratio (significance calculated using Wilcoxon rank-sum one-sided tests, parents (*n*=56) vs. replicates (*n*=174): *P*<10^-15^; parents vs. theoretical-experimental (*n*=334): *P*=1.8×10^-7^; theoretical-experimental vs. replicates: *P*<10^-15^). b) Variation of community composition across mixtures and mixture ratios. Each plot shows the Jensen-Shannon divergence (JSD) of all co-culture and theoretical communities relative to both parent communities (brown and gold) for a single mixture. Points are colored as in Fig. 1e.

**Extended Data Figure 6:**
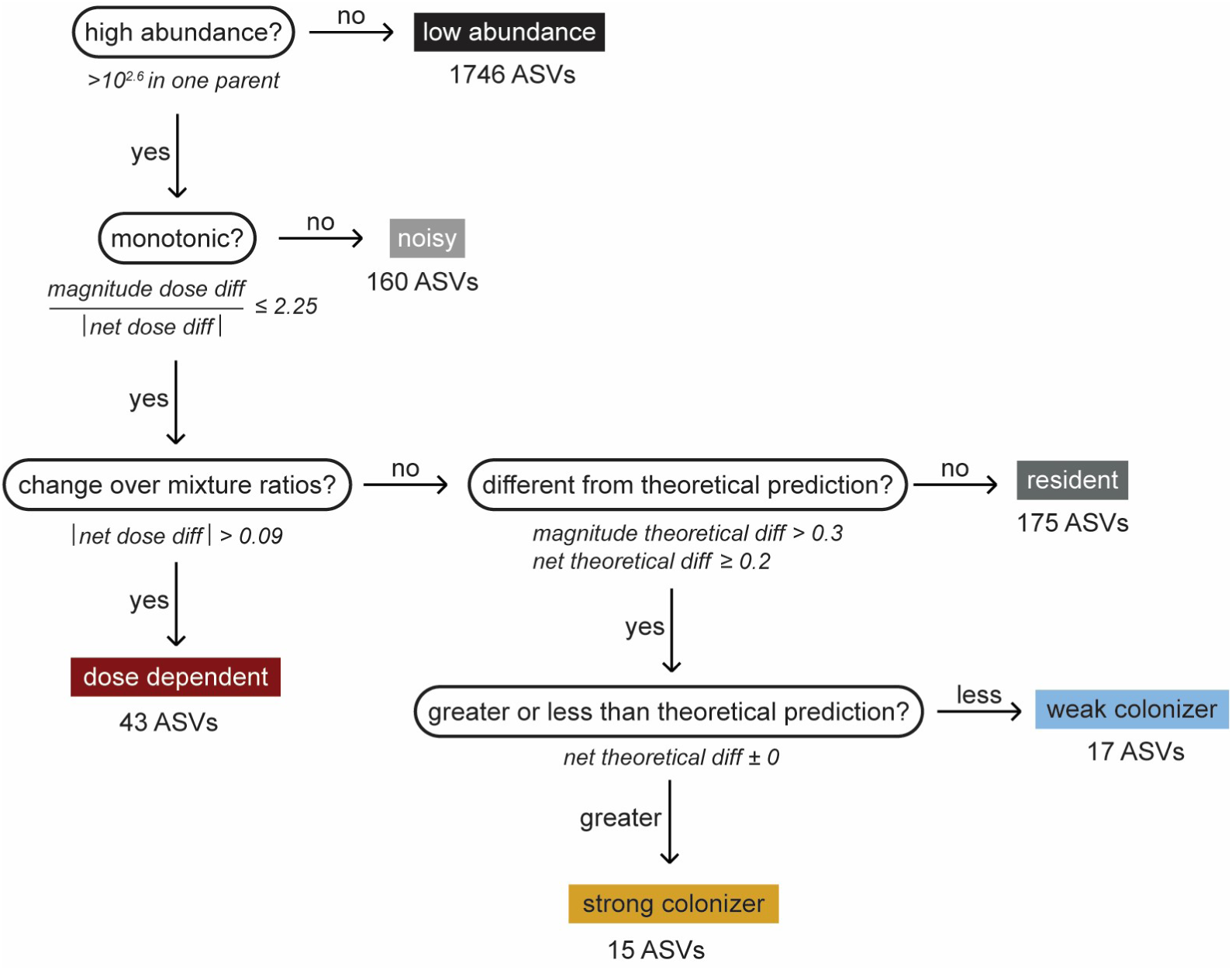
Classification scheme for ASV colonization behavior. The total number of ASVs across all mixtures is provided for each category of behavior.

**Extended Data Figure 7:**
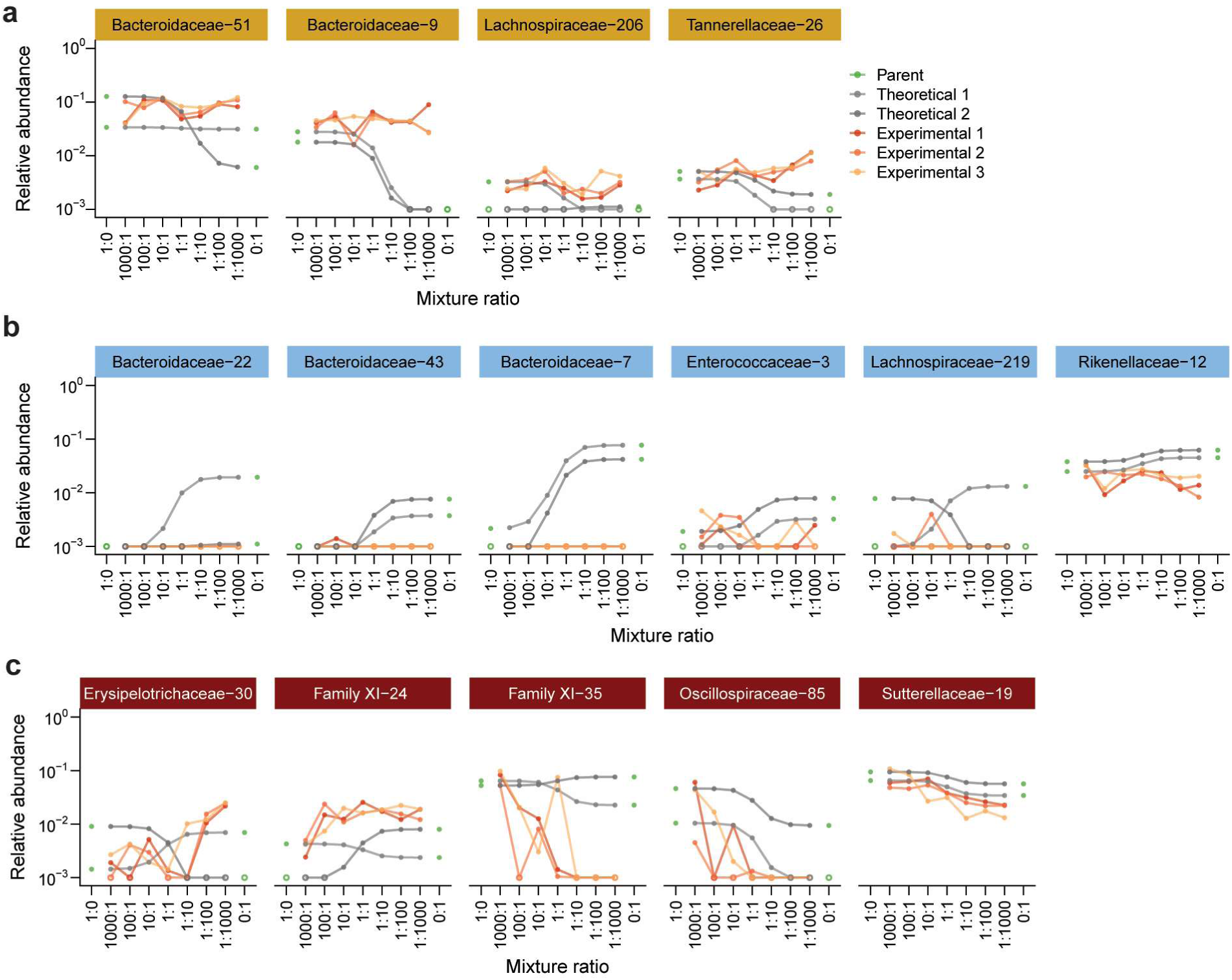
Examples of strong, weak, and dose-dependent colonizers from the C1/C2 mixture. a-c) Relative abundance of (**a**) strong, (**b**) weak, and (**c**) dose-dependent colonizers from the C1/C2 mixture. Each panel shows one ASV. Lines are colored as in Fig. 2a.

**Extended Data Figure 8:**
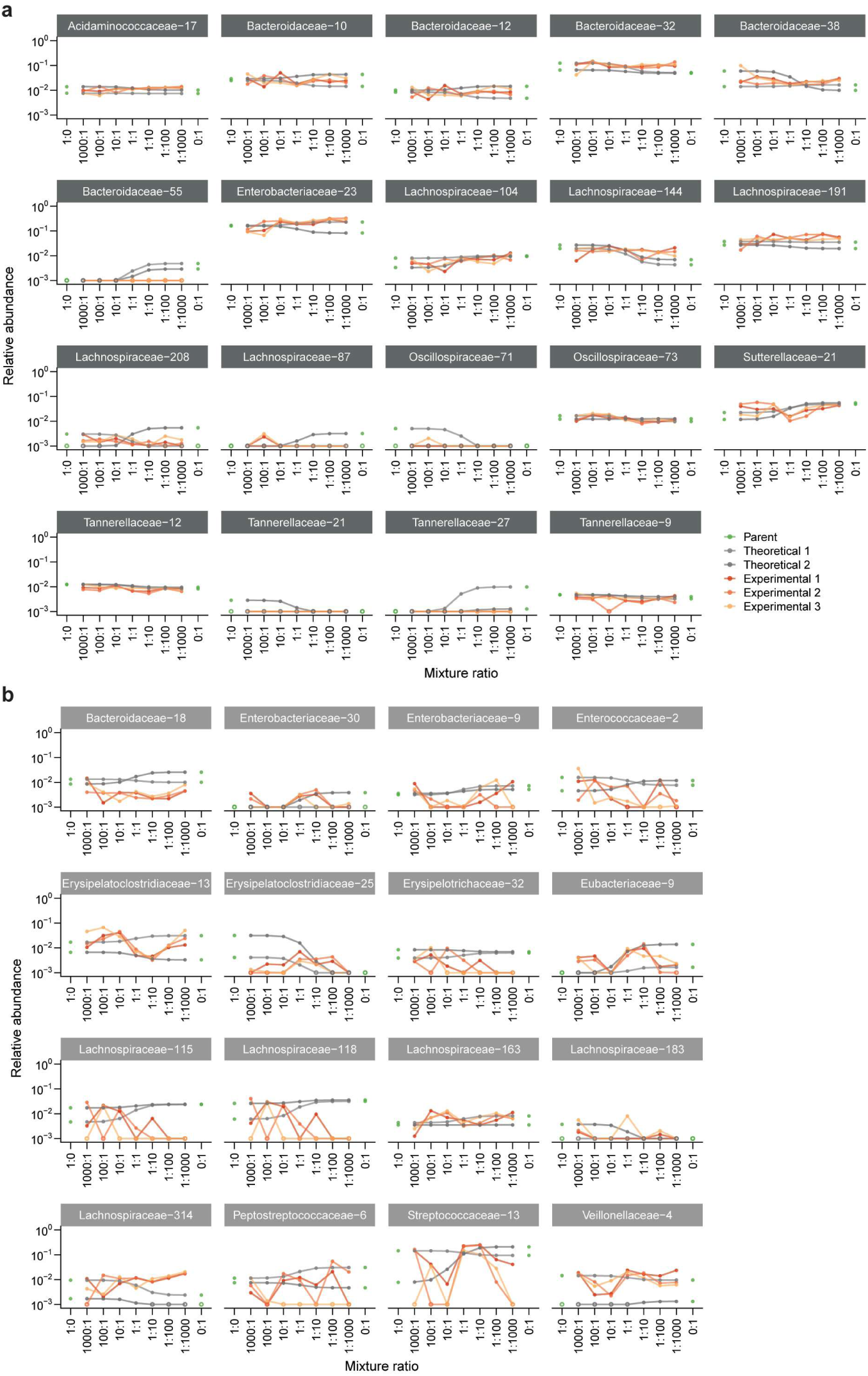
Examples of resident and noisy ASVs from the C1/C2 mixture. a,b) Relative abundance of (**a**) resident and (**b**) noisy ASVs from the C1/C2 mixture. Each panel shows one ASV. Lines are colored as in Fig. 2a.

**Extended Data Figure 9:**
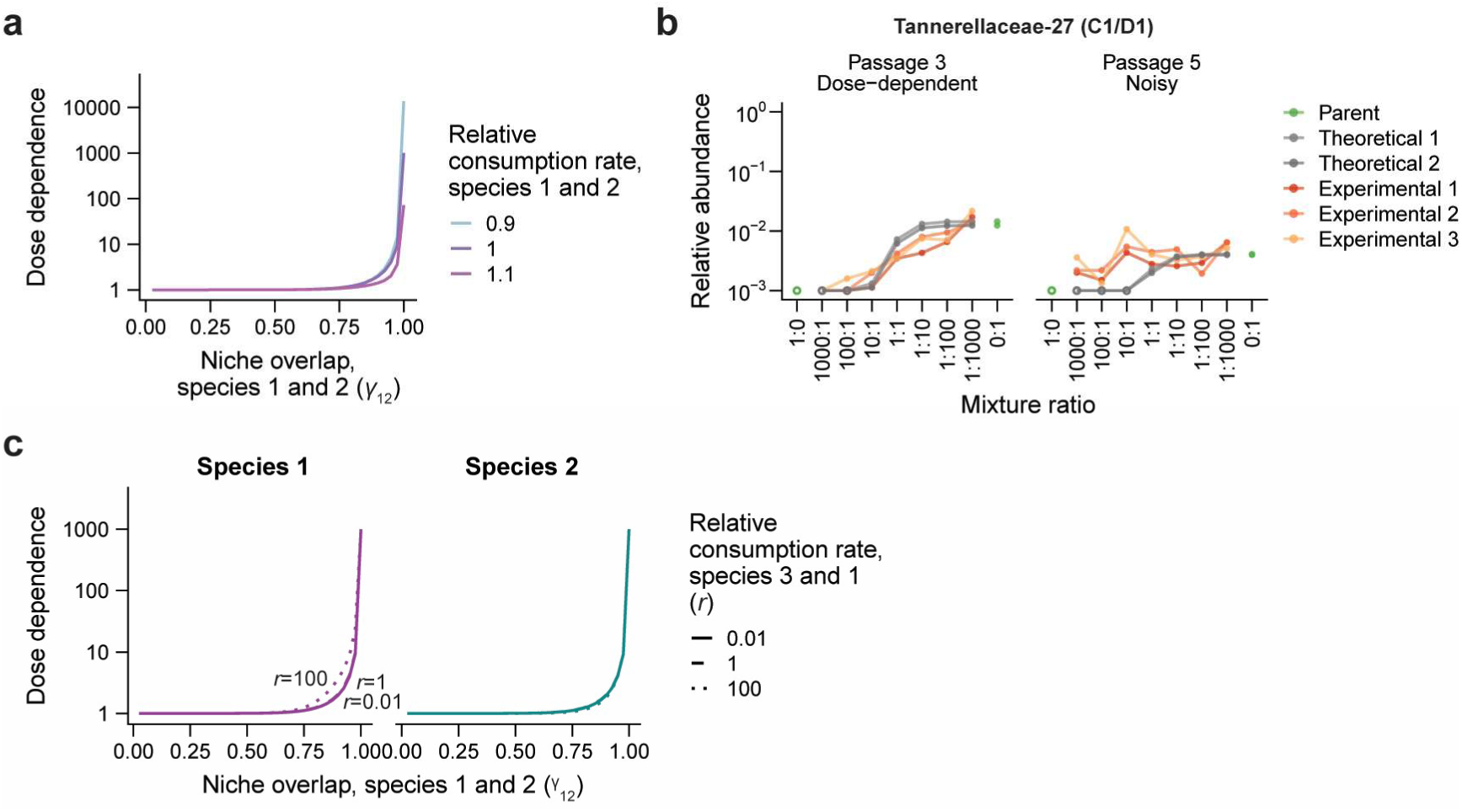
Predicted effects of niche overlap and resource consumption rates in a consumer-resource model. a) Dose dependence after five passages in a two-species consumer-resource model is robust to ∼10% variation in relative consumption rates *R*_1,12_ and *R*_2,12_ of a shared resource μ_12_. b) The dose dependence of an ASV in the Tannerellaceae family declined substantially between passages three and five. Lines are colored as in Fig. 2a. c) In our consumer-resource model, the dose dependence of species 1 after five passages increases with the relative resource consumption rate of species 3 compared with species 1 (*r*) at constant *β* = 0.5, while the dose dependence of species 2 is largely unaffected. Dose dependence is defined as in Fig. 3c.

**Extended Data Figure 10:**
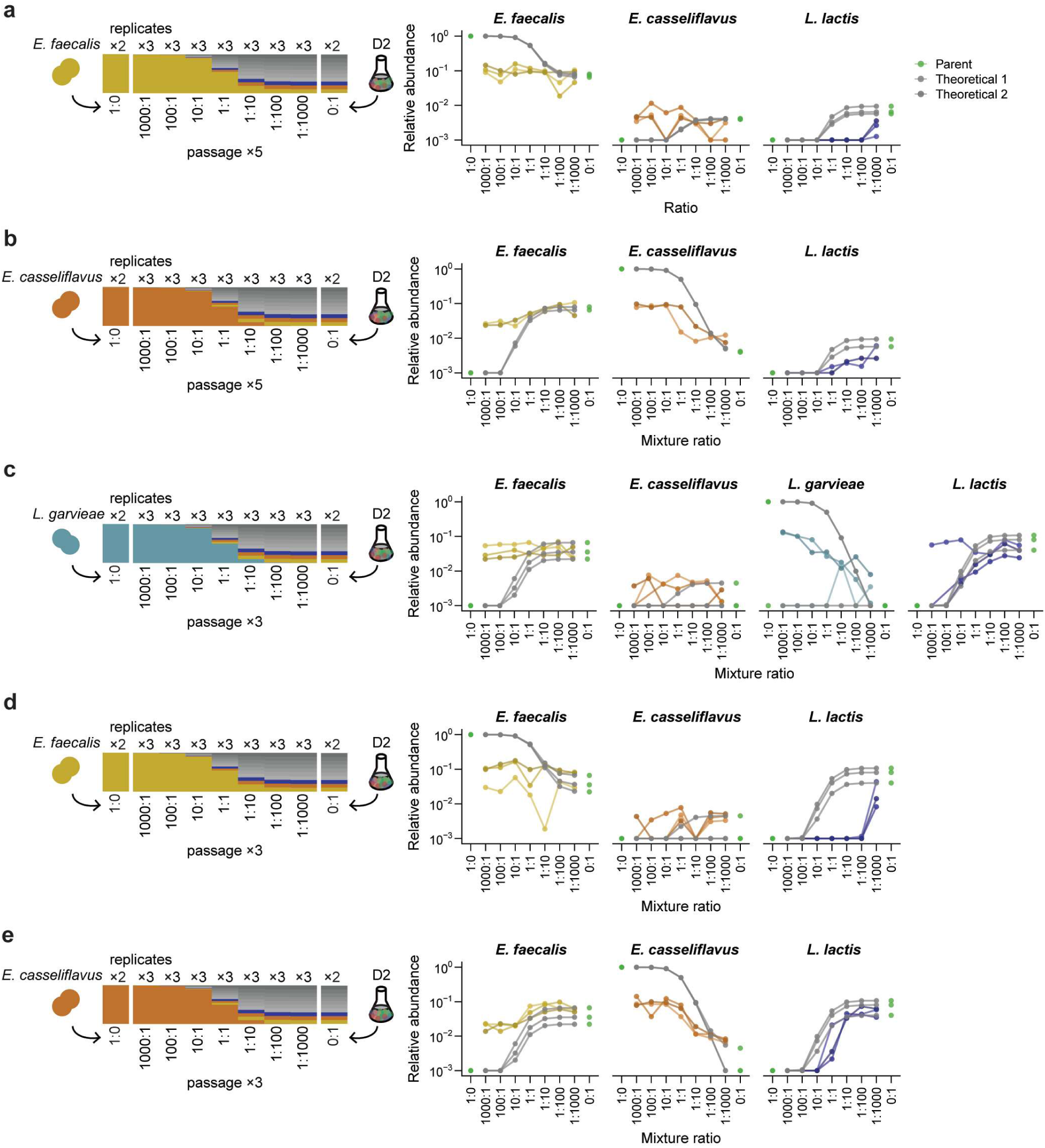
Colonization behavior in strain-community mixtures. b) Species relative abundances in the *E. faecalis*/D2 co-cultures after five passages. Only *L. lactis* was classified as dose dependent. c) Species relative abundances in the *E. casseliflavus*/D2 co-cultures after five passages. *E. faecalis*, *E. casseliflavus*, and *L. lactis* were classified as dose dependent. d) Species relative abundances in the *L. garvieae*/D2 co-cultures after three passages. *L. garvieae* and *L. lactis* were classified as dose dependent. e) Species relative abundances in the *E. faecalis*/D2 co-cultures after three passages. Only *L. lactis* was classified as dose dependent. f) Species relative abundances in the *E. casseliflavus*/D2 co-cultures after three passages. *E. faecalis*, *E. casseliflavus*, and *L. lactis* were classified as dose dependent. Lines are colored as in Fig. 2a.

**Table S1.** Sample origin of each *in vitro* community. *In vitro* communities were derived from day 0 pre-antibiotic samples collected from

| Community | Subject | Household |
| --- | --- | --- |
| A1 | XBA | XB |
| A2 | XBB | XB |
| B1 | XCA | XC |
| B2 | XCB | XC |
| C1 | XDA | XD |
| C2 | XDB | XD |
| D1 | XFA | XF |
| D2 | XFB | XF |

## Notes

### Competing Interest Statement

The authors have declared no competing interest.

### Summary of Updates

Abstract, introduction, results, and discussion sections revised to streamline results.

## References

1. Elton, C. S. The Ecology of Invasions by Animals and Plants. (Springer New York, 1958). doi:10.1007/978-1-4899-7214-9.

2. Lodge, D. M. Biological invasions: Lessons for ecology. Trends Ecol Evol 8, 133–137 (1993).

3. Williamson, M. H. & Fitter, A. The characters of successful invaders. Biol Conserv 78, 163–170 (1996).

4. Mack, R. N. et al. Biotic Invasions: Causes, Epidemiology, Global Consequences, and Control. Ecological Applications 10, 689 (2000).

5. Simberloff, D. The role of propagule pressure in biological invasions. Annu Rev Ecol Evol Syst 40, 81–102 (2009).

6. Dassonville, N., Guillaumaud, N., Piola, F., Meerts, P. & Poly, F. Niche construction by the invasive Asian knotweeds (species complex Fallopia): Impact on activity, abundance and community structure of denitrifiers and nitrifiers. Biol Invasions 13, 1115–1133 (2011).

7. Callahan, B. J., Fukami, T. & Fisher, D. S. Rapid evolution of adaptive niche construction in experimental microbial populations. Evolution (N Y) 68, 3307–3316 (2014).

8. Finlay, B. J. & Clarke, K. J. Ubiquitous dispersal of microbial species. Scientific Correspondence 400, 828 (1999).

9. Barbour, K. M., Barrón-Sandoval, A., Walters, K. E. & Martiny, J. B. H. Towards quantifying microbial dispersal in the environment. Environ Microbiol 25, 137–142 (2023).

10. Custer, G. F., Bresciani, L. & Dini-Andreote, F. Ecological and Evolutionary Implications of Microbial Dispersal. Frontiers in Microbiology vol. 13 Preprint at 10.3389/fmicb.2022.855859 (2022).

11. Shaffer, M. L. Minimum Population Sizes for Species Conservation. Bioscience 31, 131–134 (1981).

12. Lande, R. Genetics and Demography in Biological Conservation. Science (1979) 241, 1455–1460 (1985).

13. Nummi, P. Introduced Semiaquatic Birds and Mammals in Europe. in Invasive Aquatic Species of Europe. Distribution, Impacts, and Management (eds. Leppäkoski, E., Gollasch, S. & Olenin, S.) 162–172 (Springer Dordrecht, 2002).

14. Fukami, T. Historical Contingency in Community Assembly: Integrating Niches, Species Pools, and Priority Effects. Annu Rev Ecol Evol Syst 46, 1– 23 (2015).

15. Vellend, M. The Theory of Ecological Communities. (Princeton University Press, 2016).

16. Custer, G. F., Bresciani, L. & Dini-Andreote, F. Toward an integrative framework for microbial community coalescence. Trends Microbiol (2023) doi:10.1016/j.tim.2023.09.001.

17. Blackburn, T. M. et al. A proposed unified framework for biological invasions. Trends Ecol Evol 26, 333–339 (2011).

18. Hubbell, S. P. The Unified Neutral Theory of Biodiversity and Biogeography. (Princeton University Press, 2001).

19. Chesson, P. MacArthur’s Consumer-Resource Model. Theor Popul Biol 37, 26–38 (1990).

20. Lockwood, J. L., Cassey, P. & Blackburn, T. The role of propagule pressure in explaining species invasions. Trends Ecol Evol 20, 223–228 (2005).

21. Vila, J. C. C., Jones, M. L., Patel, M., Bell, T. & Rosindell, J. Uncovering the rules of microbial community invasions. Nat Ecol Evol 3, 1162–1171 (2019).

22. Ho, P.-Y., Nguyen, T. H., Sanchez, J. M., DeFelice, B. C. & Huang, K. C. Resource competition predicts assembly of in vitro gut bacterial communities. bioRxiv (2022) doi:10.1101/2022.05.30.494065.

23. Veltman, C. J., Nee, S. & Crawley, M. J. Correlates of Introduction Success in Exotic New Zealand Birds. Am Nat 147, 542–557 (1996).

24. Drake, J. M., Baggenstos, P. & Lodge, D. M. Propagule pressure and persistence in experimental populations. Biol Lett 1, 480–483 (2005).

25. Memmott, J., Craze, P. G., Harman, H. M., Syrett, P. & Fowler, S. V. The Effect of Propagule Size on the Invasion of an Alien Insect. Source: Journal of Animal Ecology 74, 50–62 (2005).

26. Acosta, F., Zamor, R. M., Najar, F. Z., Roe, B. A. & Hambright, K. D. Dynamics of an experimental microbial invasion. Proc Natl Acad Sci U S A 112, 11594–11599 (2015).

27. Pyšek, P. et al. Naturalization of central European plants in North America: Species traits, habitats, propagule pressure, residence time. Ecology 96, 762–774 (2015).

28. Chadwell, T. B. & Engelhardt, K. A. M. Effects of pre-existing submersed vegetation and propagule pressure on the invasion success of Hydrilla verticillata. Journal of Applied Ecology 45, 515–523 (2008).

29. Evans, S., Martiny, J. B. H. & Allison, S. D. Effects of dispersal and selection on stochastic assembly in microbial communities. ISME Journal 11, 176–185 (2017).

30. Albright, M. B. N., Sevanto, S., Gallegos-Graves, L. V. & Dunbar, J. Biotic Interactions Are More Important than Propagule Pressure in Microbial Community Invasions. ASM Journals mBio 11, (2020).

31. Alzate, A., Onstein, R. E., Etienne, R. S. & Bonte, D. The role of preadaptation, propagule pressure and competition in the colonization of new habitats. Oikos 129, 820–829 (2020).

32. Rojas-Botero, S., Kollmann, J. & Teixeira, L. H. Competitive trait hierarchies of native communities and invasive propagule pressure consistently predict invasion success during grassland establishment. Biol Invasions 24, 107– 122 (2022).

33. Sierocinski, P., Soria Pascual, J., Padfield, D., Salter, M. & Buckling, A. The impact of propagule pressure on whole community invasions in biomethane-producing communities. iScience 24, (2021).

34. Jones, M. L., Ramoneda, J., Rivett, D. W. & Bell, T. Biotic resistance shapes the influence of propagule pressure on invasion success in bacterial communities. Ecology 98, 1743–1749 (2017).

35. Walter, J., Maldonado-Gómez, M. X. & Martínez, I. To engraft or not to engraft: an ecological framework for gut microbiome modulation with live microbes. Curr Opin Biotechnol 49, 129–139 (2018).

36. Suez, J., Zmora, N., Segal, E. & Elinav, E. The pros, cons, and many unknowns of probiotics. Nature Medicine vol. 25 716–729 Preprint at 10.1038/s41591-019-0439-x (2019).

37. Gianotti, L. et al. A randomized double-blind trial on perioperative administration of probiotics in colorectal cancer patients. World J Gastroenterol 16, 167–175 (2010).

38. Maldonado-Gómez, M. X. et al. Stable Engraftment of Bifidobacterium longum AH1206 in the Human Gut Depends on Individualized Features of the Resident Microbiome. Cell Host Microbe 20, 515–526 (2016).

39. Ouwehand, A. C. A review of dose-responses of probiotics in human studies. Benef Microbes 8, 143–151 (2017).

40. Zmora, N. et al. Personalized Gut Mucosal Colonization Resistance to Empiric Probiotics Is Associated with Unique Host and Microbiome Features. Cell 174, 1388–1405.e21 (2018).

41. Li, S. S. et al. Durable coexistence of donor and recipient strains after fecal microbiota transplantation. Science (1979) 352, 586–589 (2016).

42. Schmidt, T. S. B. et al. Drivers and determinants of strain dynamics following fecal microbiota transplantation. Nat Med 28, 1902–1912 (2022).

43. Jessup, C. M. et al. Big questions, small worlds: Microbial model systems in ecology. Trends Ecol Evol 19, 189–197 (2004).

44. Aranda-Díaz, A., et al. Assembly of gut-derived bacterial communities follows ‘early-bird’ resource utilization dynamics. bioRxiv (2023) doi:10.1101/2023.01.13.523996.

45. Javdan, B. et al. Personalized Mapping of Drug Metabolism by the Human Gut Microbiome. Cell 181, 1661–1679.e22 (2020).

46. Aranda-Díaz, A. et al. Establishment and characterization of stable, diverse, fecal-derived in vitro microbial communities that model the intestinal microbiota. Cell Host Microbe 30, 260–272.e5 (2022).

47. Celis, A. I., Relman, D. A. & Huang, K. C. The impact of iron and heme availability on the healthy human gut microbiome in vivo and in vitro. Cell Chem Biol 30, 110–126.e3 (2023).

48. Rillig, M. C. et al. Interchange of entire communities: Microbial community coalescence. Trends Ecol Evol 30, 470–476 (2015).

49. Posfai, A., Taillefumier, T. & Wingreen, N. S. Metabolic Trade-Offs Promote Diversity in a Model Ecosystem. Phys Rev Lett 118, (2017).

50. Abreu, C. I., Andersen Woltz, V. L., Friedman, J. & Gore, J. Microbial communities display alternative stable states in a fluctuating environment. PLoS Comput Biol 16, (2020).

51. Pereira, F. C. & Berry, D. Microbial nutrient niches in the gut. Environ Microbiol 19, 1366–1378 (2017).

52. Sprockett, D., Fukami, T. & Relman, D. A. Role of priority effects in the early-life assembly of the gut microbiota. Nat Rev Gastroenterol Hepatol 15, 197–205 (2018).

53. Wootton, J. T. Field parameterization and experimental test of the neutral theory of biodiversity. Nature 433, 309–312 (2005).

54. D’Andrea, R., Gibbs, T. & O’Dwyer, J. P. Emergent neutrality in consumer-resource dynamics. PLoS Comput Biol 16, (2020).

55. Grilli, J., Barabás, G., Michalska-Smith, M. J. & Allesina, S. Higher-order interactions stabilize dynamics in competitive network models. Nature 548, 210–213 (2017).

56. Morin, M. A., Morrison, A. J., Harms, M. J. & Dutton, R. J. Higher-order interactions shape microbial interactions as microbial community complexity increases. Sci Rep 12, (2022).

57. Ludington, W. B. Higher-order microbiome interactions and how to find them. Trends Microbiol 30, 618–621 (2022).

58. Chang, C.-Y., Bajić, D., Vila, J. C. C., Estrela, S. & Sanchez, A. Emergent coexistence in multispecies microbial communities. Science (1979) 381, (2023).

59. Fukami, T. Assembly history interacts with ecosystem size to influence species diversity. Ecology 85, 3234–3242 (2004).

60. Fukami, T. & Nakajima, M. Community assembly: Alternative stable states or alternative transient states? Ecol Lett 14, 973–984 (2011).

61. Song, C., Fukami, T. & Saavedra, S. Untangling the complexity of priority effects in multispecies communities. Ecol Lett 24, 2301–2313 (2021).

62. Mallon, C. A. et al. The impact of failure: Unsuccessful bacterial invasions steer the soil microbial community away from the invader’s niche. ISME Journal 12, 728–741 (2018).

63. Warren, P. H., Law, R. & Weatherby, A. J. Mapping the Assembly of Protist Communities in Microcosms. Ecology 84, 1001–1011 (2003).

64. Amor, D. R., Ratzke, C. & Gore, J. Transient invaders can induce shifts between alternative stable states of microbial communities. Sci Adv 6, (2020).

65. Chappell, C. R. et al. Wide-ranging consequences of priority effects governed by an overarching factor. Elife 11, (2022).

66. Simberloff, D. & Von Holle, B. Positive interactions of nonindigenous species: invasional meltdown? Biological Invasions vol. 1 (1999).

67. Miller, T. E., TerHorst, C. P. & Burns, J. H. The ghost of competition present. American Naturalist 173, 347–353 (2009).

68. Olito, C. & Fukami, T. Long-term effects of predator arrival timing on prey community succession. American Naturalist 173, 354–362 (2009).

69. O’Callaghan, M., Ballard, R. A. & Wright, D. Soil microbial inoculants for sustainable agriculture: Limitations and opportunities. Soil Use Manag 38, 1340–1369 (2022).

70. Pudlo, N. A., et al. Phenotypic and Genomic Diversification in Complex Carbohydrate-Degrading Human Gut Bacteria. mSystems 7, (2022).

71. Han, S. et al. A metabolomics pipeline for the mechanistic interrogation of the gut microbiome. Nature 595, 415–420 (2021).

72. Abrams, P. The Theory of Limiting Similarity. Source: Annual Review of Ecology and Systematics 14, 359–376 (1983).

73. Connors, B. M., et al. Model-guided design of the diversity of a synthetic human gut community. bioRxiv (2022) doi:10.1101/2022.03.14.484355.

74. Martiny, J. B. H., Jones, S. E., Lennon, J. T. & Martiny, A. C. Microbiomes in light of traits: A phylogenetic perspective. Science vol. 350 Preprint at 10.1126/science.aac9323 (2015).

75. Xue, K. S., et al. Prolonged delays in human microbiota transmission after a controlled antibiotic perturbation. bioRxiv (2023) doi:10.1101/2023.09.26.559480.

76. Apprill, A., Mcnally, S., Parsons, R. & Weber, L. Minor revision to V4 region SSU rRNA 806R gene primer greatly increases detection of SAR11 bacterioplankton. Aquatic Microbial Ecology 75, 129–137 (2015).

77. Parada, A. E., Needham, D. M. & Fuhrman, J. A. Every base matters: Assessing small subunit rRNA primers for marine microbiomes with mock communities, time series and global field samples. Environ Microbiol 18, 1403–1414 (2016).

78. Smith, T., Heger, A. & Sudbery, I. UMI-tools: Modeling sequencing errors in Unique Molecular Identifiers to improve quantification accuracy. Genome Res 27, 491–499 (2017).

79. Martin, M. Cutadapt removes adapter sequences from high-throughput sequencing reads. EMBnet J 17, 10–12 (2011).

80. Callahan, B. J. et al. DADA2: High-resolution sample inference from Illumina amplicon data. Nat Methods 13, 581–583 (2016).

81. Quast, C. et al. The SILVA ribosomal RNA gene database project: Improved data processing and web-based tools. Nucleic Acids Res 41, (2013).

82. Shiver, A. L., Culver, R., Deutschbauer, A. M. & Huang, K. C. Rapid ordering of barcoded transposon insertion libraries of anaerobic bacteria. Nat Protoc 16, 3049–3071 (2021).

83. Cermak, N., Datta, M. Sen & Conwill, A. Rapid, Inexpensive Measurement of Synthetic Bacterial Community Composition by Sanger Sequencing of Amplicon Mixtures. iScience 23, (2020).

84. Blazanin, M. gcplyr: an R package for microbial growth curve data analysis. bioRxiv (2023) doi:10.1101/2023.04.30.538883.

